# Genetics of Latin American Diversity (GLAD) Project: insights into population genetics and association studies in recently admixed groups in the Americas

**DOI:** 10.1101/2023.01.07.522490

**Authors:** Victor Borda, Douglas P. Loesch, Bing Guo, Roland Laboulaye, Diego Veliz-Otani, Jennifer N. French-Kwawu, Thiago Peixoto Leal, Stephanie M. Gogarten, Sunday Ikpe, Mateus H. Gouveia, Marla Mendes, Gonçalo R. Abecasis, Isabela Alvim, Carlos E. Arboleda-Bustos, Gonzalo Arboleda, Humberto Arboleda, Mauricio L. Barreto, Lucas Barwick, Marcos A. Bezzera, John Blangero, Vanderci Borges, Omar Caceres, Jianwen Cai, Pedro Chana-Cuevas, Zhanghua Chen, Brian Custer, Michael Dean, Carla Dinardo, Igor Domingos, Ravindranath Duggirala, Elena Dieguez, Willian Fernandez, Henrique B. Ferraz, Frank D. Gilliland, Heinner Guio, Bernardo Horta, Joanne E. Curran, Jill M. Johnsen, Robert C. Kaplan, Shannon Kelly, Eimear E. Kenny, Barbara A. Konkle, Charles Kooperberg, Andres Lescano, M. Fernanda Lima-Costa, Ruth J. F. Loos, Ani Manichaikul, Deborah A. Meyers, Michel S. Naslavsky, Deborah A. Nickerson, Kari E. North, Carlos Padilla, Michael Preuss, Victor Raggio, Alexander P. Reiner, Stephen S. Rich, Carlos R. Rieder, Michiel Rienstra, Jerome I. Rotter, Tatjana Rundek, Ralph L. Sacco, Cesar Sanchez, Vijay G. Sankaran, Bruno Lopes Santos-Lobato, Artur Francisco Schumacher-Schuh, Marilia O. Scliar, Edwin K. Silverman, Tamar Sofer, Jessica Lasky-Su, Vitor Tumas, Scott T. Weiss, Latin American Research Consortium on the Genetics of Parkinson’s Disease (LARGE-PD), NINDS Stroke Genetics Network (SiGN) Consortium, TOPMed Population Genetics Working Group, Ignacio F. Mata, Ryan D. Hernandez, Eduardo Tarazona-Santos, Timothy D. O’Connor

## Abstract

Latin America is underrepresented in genetic studies, which can exacerbate disparities in personalized genomic medicine. However, genetic data of thousands of Latin Americans are already publicly available, but require a bureaucratic maze to navigate all the data access and consenting issues. We present the Genetics of Latin American Diversity (GLAD) Project, a platform that compiles genome-wide information of 54,077 Latin Americans from 39 studies representing 45 geographical regions. Through GLAD, we identified heterogeneous ancestry composition and recent gene-flow across the Americas. Also, we developed a simulated-annealing-based algorithm to match the genetic background of external samples to our database and share summary statistics without transferring individual-level data. Finally, we demonstrate the potential of GLAD as a critical resource for evaluating statistical genetic softwares in the presence of admixture. By making this resource available, we promote genomic research in Latin Americans and contribute to the promises of personalized medicine to more people.

## Introduction

Latin Americans/Latinos/Latinx/Latine, or Hispanics, as an ethnic label, represent a set of populations across the Americas characterized by admixture between populations from many parts of the world with distinct ancestry compositions ^1^. As such, treating Latin Americans as a single group is an over-simplification that may limit opportunities to improve health and clinical treatment. Latin Americans comprise 656 million people (8.5% of the world’s population)^2^. In the United States, Latin Americans represent 18% of the population and are the fastest-growing demographic^3^. Unfortunately, these populations remain understudied and underserved in biomedical research and are at risk of being left behind by the precision medicine revolution. For example, Latin Americans only represent about 0.23% of participants in genome-wide association studies (GWAS) performed^4^. Several important efforts have been made to understand Latin American (**LAm**) genetic history and to identify genetic variants associated with complex traits ^5–26^. However, most of these samples are thinly spread across many projects with few initiatives (e.g., the Mexico City Prospective Study^27^) to obtain the 100K+ individuals necessary to have statistical power comparable to other population groups (e.g., Europeans^28^ and East Asians^29,30^).

To remedy the under-representation of Latin Americans in genomic studies, we have created the Genetics of Latin American Diversity database (GLADdb), a resource to infer fine-scale patterns of population structure across the Americas and boost statistical power for the discovery of genetic factors contributing to LAm health and disease. By gleaning LAm individuals through dbGaP and whole genome sequencing projects across the Americas, we gathered over 54,000 unrelated individuals, either genotyped and imputed, or sequenced, from ten countries (**Figure 1A**) spanning 45 geographical groups (**Table S1** and **Table S2**). These group labels reflect administrative division level (e.g., country, state, or city level information) when available. Using GLADdb, we addressed two major goals regarding LAm genomics: (i) in population genetics: to identify recent fine-scale patterns of distant relatedness and differentiation along the Americas, providing insights into regions with genetic underrepresentation, and (ii) in genetic epidemiology at two levels: (a) by developing a web tool for matching the genetic background of GLADdb individuals with external pools of samples providing additional power to discover genotype-phenotype associations and (b) by demonstrating how GLADdb can be utilized for testing statistical genetic software in diverse LAm cohorts.

**Figure 1.**
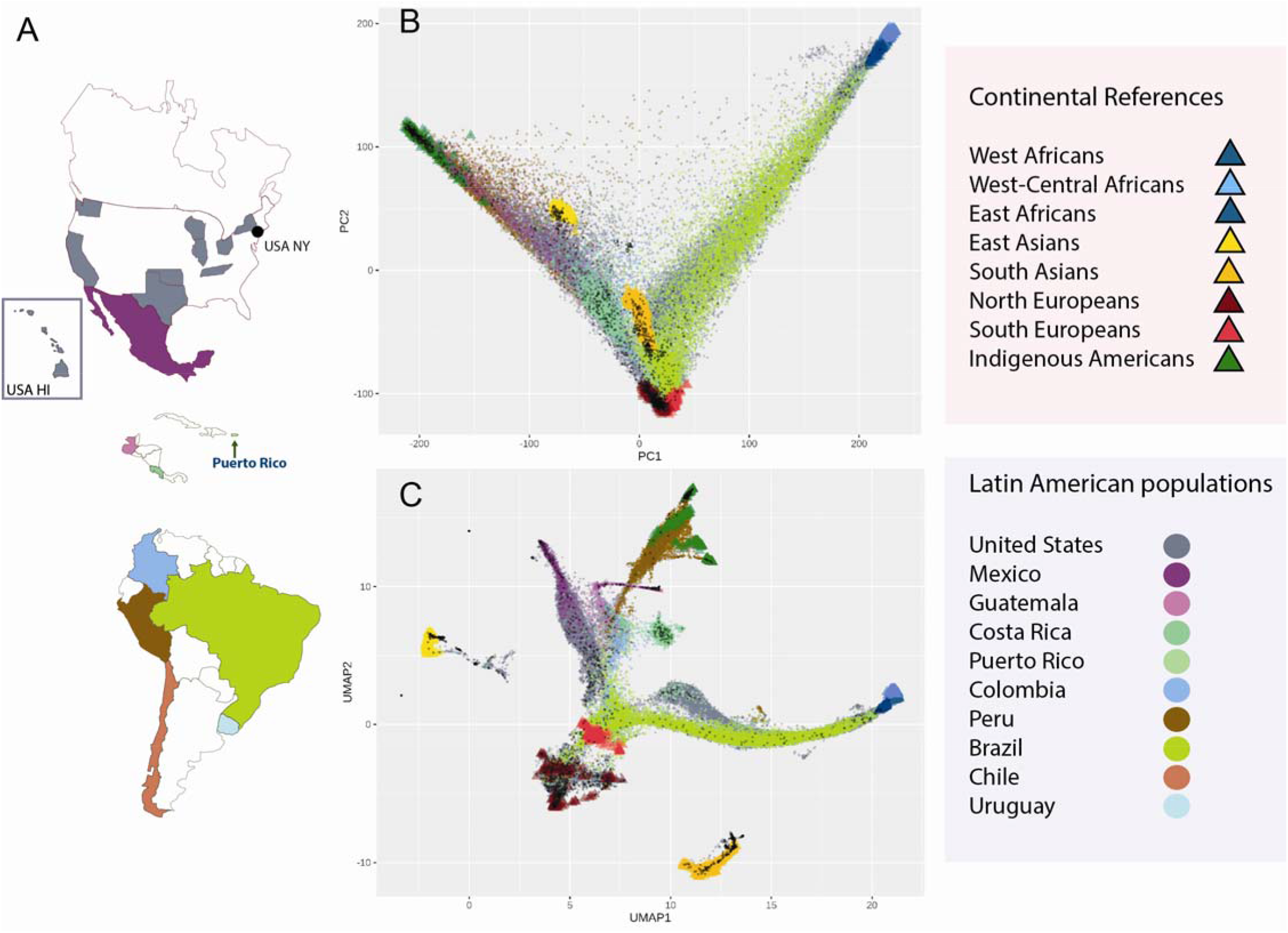
Dimensionality reduction of genetic data for more than 52K unrelated Latin Americans from the GLAD database. A) Geographical distribution of GLADdb cohorts. B) Principal Component Analysis of the entire dataset based on high-quality imputed SNPs (r^2^ > 0.9) showing the sampling spread of Latin Americans. C) Uniform Manifold Approximation and Projection (UMAP) of the first 10 PCs showing clusters of different population groups.

We start by exploring distant genetic relatedness among LAm countries. Several studies have focused on determining the sources and timing for admixture events that led to the current genetic composition in some LAm countries ^6,11–13,16,31–33^. However, understanding LAm genetic diversity goes beyond the initial continental admixture and involves bottlenecks, founder effects, and migration into and along the Americas, especially as it relates to fine-scale population structure within the continental sources (i.e., Indigenous American, European and African groups). We explored population structure and recent migration among LAm regions by analyzing allelic frequencies and identity-by-descent (IBD) sharing.

We then address issues about data availability when performing large-scale analyses in LAm populations. Many association analyses in LAm populations have smaller sample sizes than similar studies in Europeans and other populations. Data, even when publicly available, is often prohibitively restrictive for investigators to access because of quality control efforts and data curation, in addition to the bureaucratic maze typically required to obtain the data ^34^. Artomov et al. ^35^ showed that with a large control cohort, a matching procedure, which is the identification of individuals with similar genetic backgrounds with external data, and sharing of their summary statistics (e.g., allele counts), is possible without the transfer of individual-level data. The matching procedure is designed to guard against genetic control inflation and reduce spurious associations due to population structure. Given Artomov’s approach was developed on European ancestry individuals, we adapt this idea to the complex ancestral composition of LAm individuals. We devised an enhanced matching algorithm to explore the principal component space derived from our diverse GLADdb cohorts, into which we project external samples and match them to GLADdb individuals using ancestral background summary statistics. From the selected GLADdb individuals, we will generate and return summary statistics of genome-wide genotype frequencies and aggregate local ancestry composition to increase the sample size and power of the end-user study. Since GLADdb consists of both cases and controls for different phenotypes, we will also use phenotype filters to select individuals useful as controls. We implemented all these features through an interactive web portal (glad.igs.umaryland.edu).

Finally, we demonstrate the potential of GLADdb as a critical resource for evaluating the performance of statistical genetic software in the presence of admixture. We do so by comparing three polygenic risk score (PRS) algorithms for estimating PRS in admixed individuals in a scenario where the ancestries corresponding to the GWAS summary statistics do not match the target cohort. PRS, the linear summation of risk variants weighted by their GWAS effect size, are highly impacted by the European-ancestry bias underlying much of the available GWAS data, and their transferability across populations remains a critical limitation of the approach ^36,37^. GLADdb is uniquely situated to support methods development efforts that help ensure cross-population transferability of statistical genetic applications.

## Results

### Data Description and QC

Our main workflow is described in **Figure S1** and **Supplementary methods**. Briefly, we have explored over 268K samples by gathering data from 39 dbGaP cohorts and other WGS projects that include US Hispanics / LAm individuals ^5–8,13,38^ (**Table S1**). As inclusion criteria, we gathered individuals self-described as “Latino” or “Hispanic” and ADMIXTURE-defined individuals. This latter criterion was applied to identify possible LAm individuals using ADMIXTURE analysis^39^, keeping any individuals with more than 2% Indigenous American (IA) ancestry (See Methods). For genotyped cohorts (**Table S1**), we imputed all self-described (GLAD-SD, n=25,627) and ADMIXTURE-defined (GLAD-AD, n=17,642) individuals within each cohort using the TOPMed Imputation server^40^. After imputation QC, we kept 42,539 individuals that were combined into a single dataset with sequencing data from TOPMed Project^5^ (27,088 individuals) and 1000 Genomes Project^38^ (345 individuals) with 9,121,629 overlapping variants with an imputation r^2^ > 0.3 across all datasets. For all analyses here, we kept overlapping variants with imputation r^2^ > 0.9 in each dataset before merging. The final merged dataset with r^2^ > 0.9 for analysis contains 3,248,494 biallelic variants. Finally, to remove the family structure in GLADdb, we inferred kinship coefficients using IBD segments on the complete dataset, keeping 54,077 unrelated individuals (See Methods).

### Continental Population Structure of GLADdb

Using 54K unrelated samples and ancestry-reference groups (**Table S3**), we explored the patterns of diversity and differentiation throughout the Americas using principal component analysis (PCA), uniform manifold approximation and projection (UMAP), and ADMIXTURE analyses (**Figure 1B and C, Figure S2-S5**). Both results highlighted some important points. First, the samples cluster according to ancestry and not technology or other batch effects (**Figures S3 and S4**). Notably, GLAD-AD individuals cluster well with other GLAD-SD individuals validating our inclusion criteria (**Figure S4A and B**). By coupling UMAP and ADMIXTURE results, we reaffirm the heterogeneous ancestry distribution of LAm individuals, with some groups showing predominantly IA ancestry (Peru, Mexico, and Guatemala) and others showing majority admixture between European and African ancestries (USA and Brazil) (**Figure 1C** and **Figure S5**). Regarding sample sizes, the best-represented regions in GLADdb included Brazil, Central America, Mexico, Peru, and the United States.

### Levels of genetic diversity within Latin American groups

Although our population structure analyses identified a wide diversity of LAm groups, these groups originated from continental progenitors that suffered a significant drop in effective population size during the colonial period of the Americas^41–43^. This resulted in a higher level of consanguinity and enrichment of long runs of homozygosity observed in some LAm groups (e.g., CLM and PEL from 1000 Genomes Project) compared to Finnish^42^, a population notably shaped by a strong founder effect. Based on demographic information available for the cohorts, we organized GLAD-SD individuals into 45 self-described LAm groups, consistent with geographic labels based on administrative division level (e.g., country, state, or city level information) (**Table S2**). In addition, we included 12 IA populations from the Peruvian Genome project as well as 5 European (EUR) and 5 African (AFR) populations from the 1000 Genomes Project (See Methods).

We explored the levels of diversity in each group by inferring runs of homozygosity^44^ (ROH) (**Figure 2**, See Methods). As expected, individuals from Africa showed lower values for total ROH compared to individuals from Europe and Indigenous groups from Peru. Analogously, LAm groups with higher proportions of African ancestry (e.g., Peru-Ica and Northeast Brazilian regions) tend to have the lowest total ROH. Furthermore, taking advantage of the detailed sample representation for 13 Peruvian and 12 Brazilian regions, we determined the correlation between average genome-wide ancestry proportions (**Table S2**) and the median total ROH for each population. We observed a positive correlation between the average Indigenous American (*r*=0.81, *p-value* = 0.00246) and European (*r*=0.88, *p-value* = 1.12 x 10^-4^) ancestries with a higher density of ROH in Peruvians and Brazilians, respectively. Interestingly, both correlations follow a North to South line.

**Figure 2.**
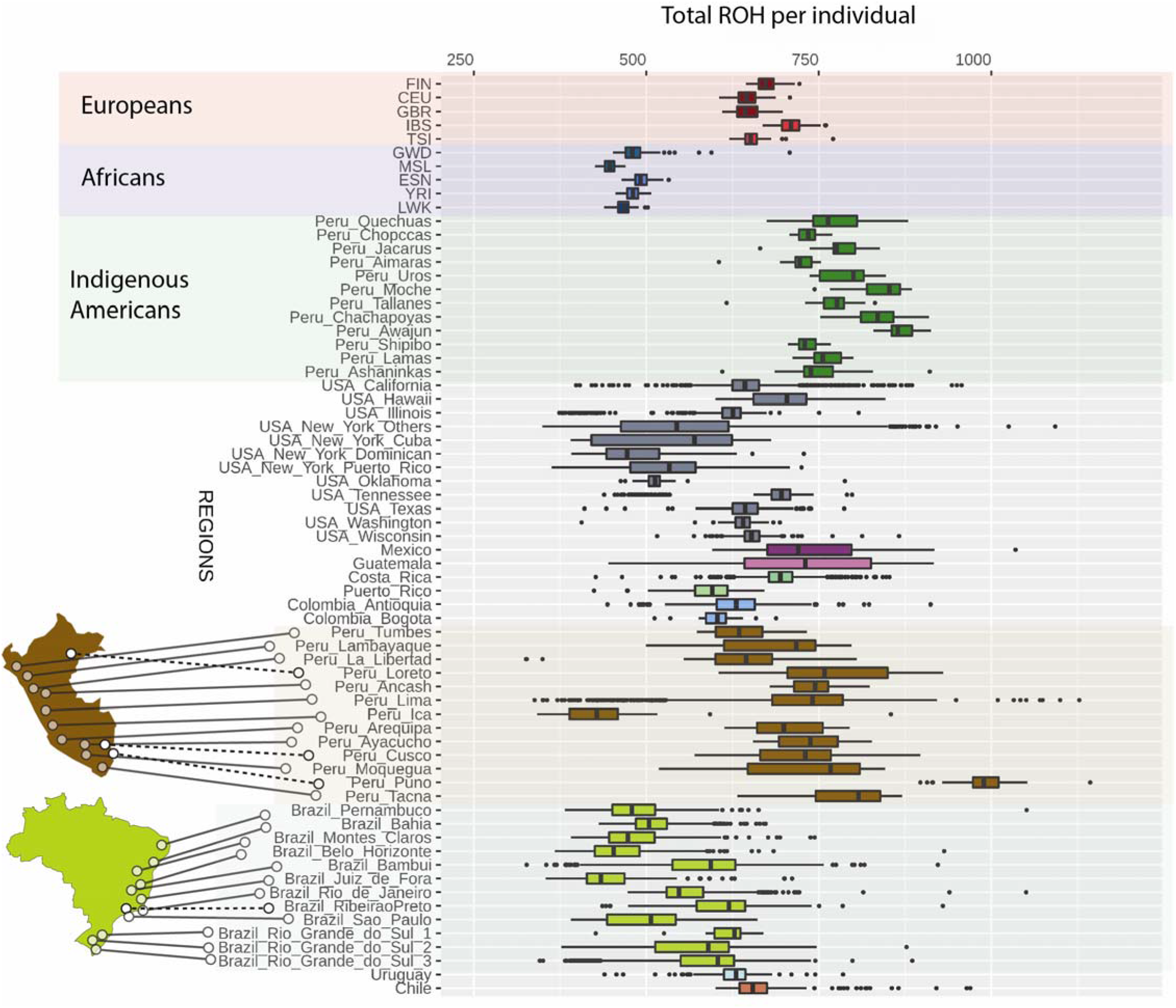
Distribution of Genome-wide amount of Runs of Homozygosity for Latin American groups and Reference populations included in GLADdb. The upper part of the plot shows continental reference populations—the lower part details the distribution in Peru and Brazil. Populations are sorted in a North-to-South pattern.

### Fine-scale population structure revealed by IBD network

To obtain a fine-scale picture of population structure among LAm groups, we built a sample-pair genome-wide total IBD matrix using all IBD segments > 5cM shared in our 54K dataset. Clusters in this matrix are mainly consistent with geographic labels, with strong intra-cluster sharing among individuals from Puerto Rico, Dominican Republic, and Costa Rica (**Figure 3**). Given the sample size and genetic diversity, finer-scale population structure is observable in clusters representing the USA/Mexico, Peru, and Brazil. To reveal the substructure, we employed an IBD network-based community detection algorithm to further analyze relatedness patterns. We selected the top 20 IBD-network-based communities that accumulated 69% of GLADdb individuals (other communities each have less than 270 individuals). Each of these communities (labeled as CA1 to 20 and ordered from largest to smallest) showed enrichment of individuals from a particular country, such as Costa Rica (99.6%, IBD community CA5), Puerto Rico (98%, IBD community CA2), Dominican Republic (95.0%, CA4), Cuba (89.8%, CA6), Colombia (89.4%, CA13), and Chile (84%, CA19) (**Figure 4**). In contrast, individuals from Mexico, Peru, and Brazil were grouped in several communities (Mexico: 7, Brazil: 5, Peru: 13 communities). These communities were represented by individuals from a particular region, reflecting the extensive sampling performed in these countries (**Figure 4**).

**Figure 3.**
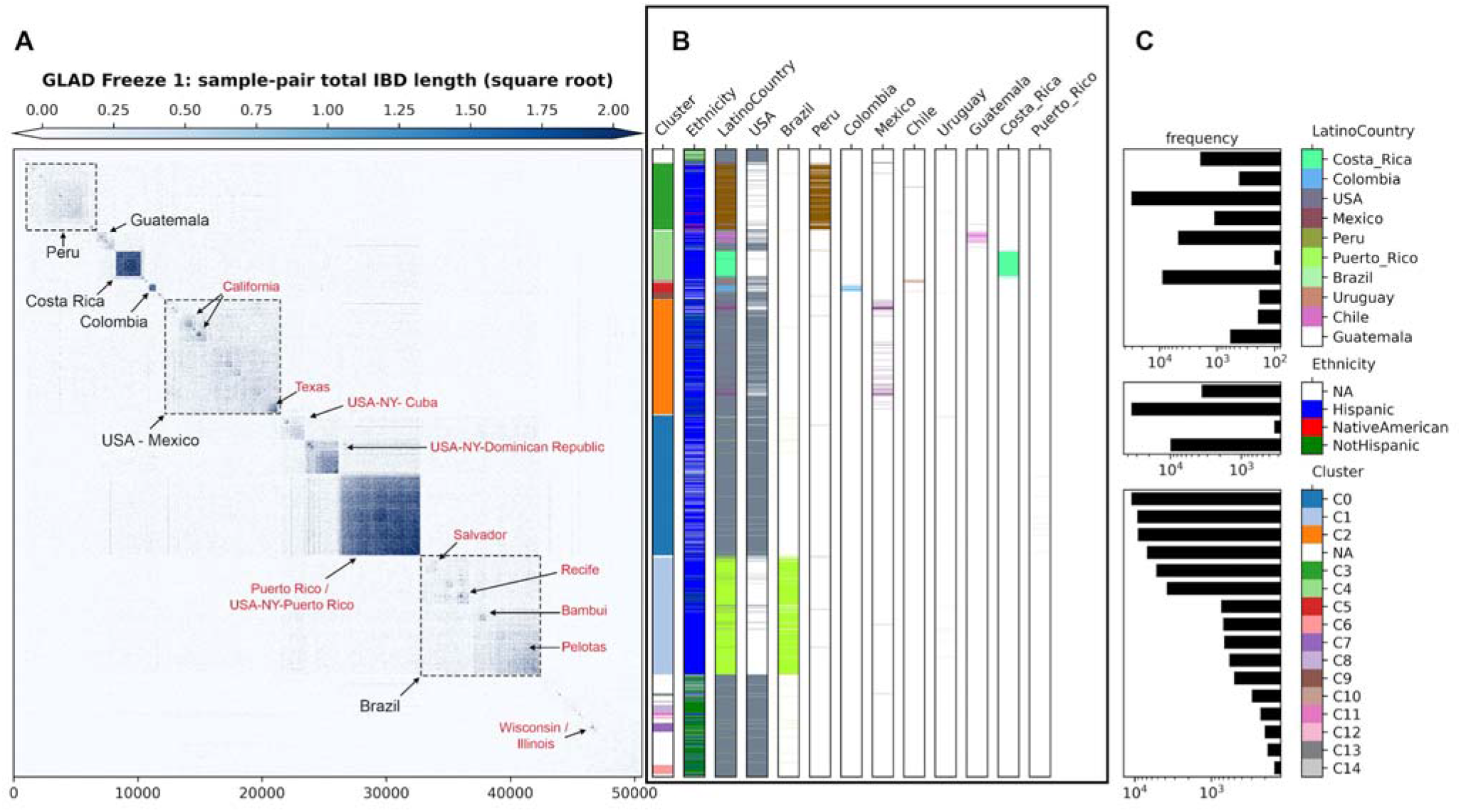
Clustering of total IBD matrix of unrelated individuals from GLADdb. A) Heatmap of the square root of sample-pair total IBD. Annotations within the heatmap represent the most enriched demographic labels in the indicated blocks. Labels with “USA-NY-country” correspond to self-described US-Hispanic living in New York with a specific country of origin. B) Individual-level labels of agglomerative cluster assignment (1st column), ethnicity (2nd), and sampling Country (combined:3rd and separated: 4th onward). C) Frequency of labels (log scale) and color keys.

**Figure 4.**
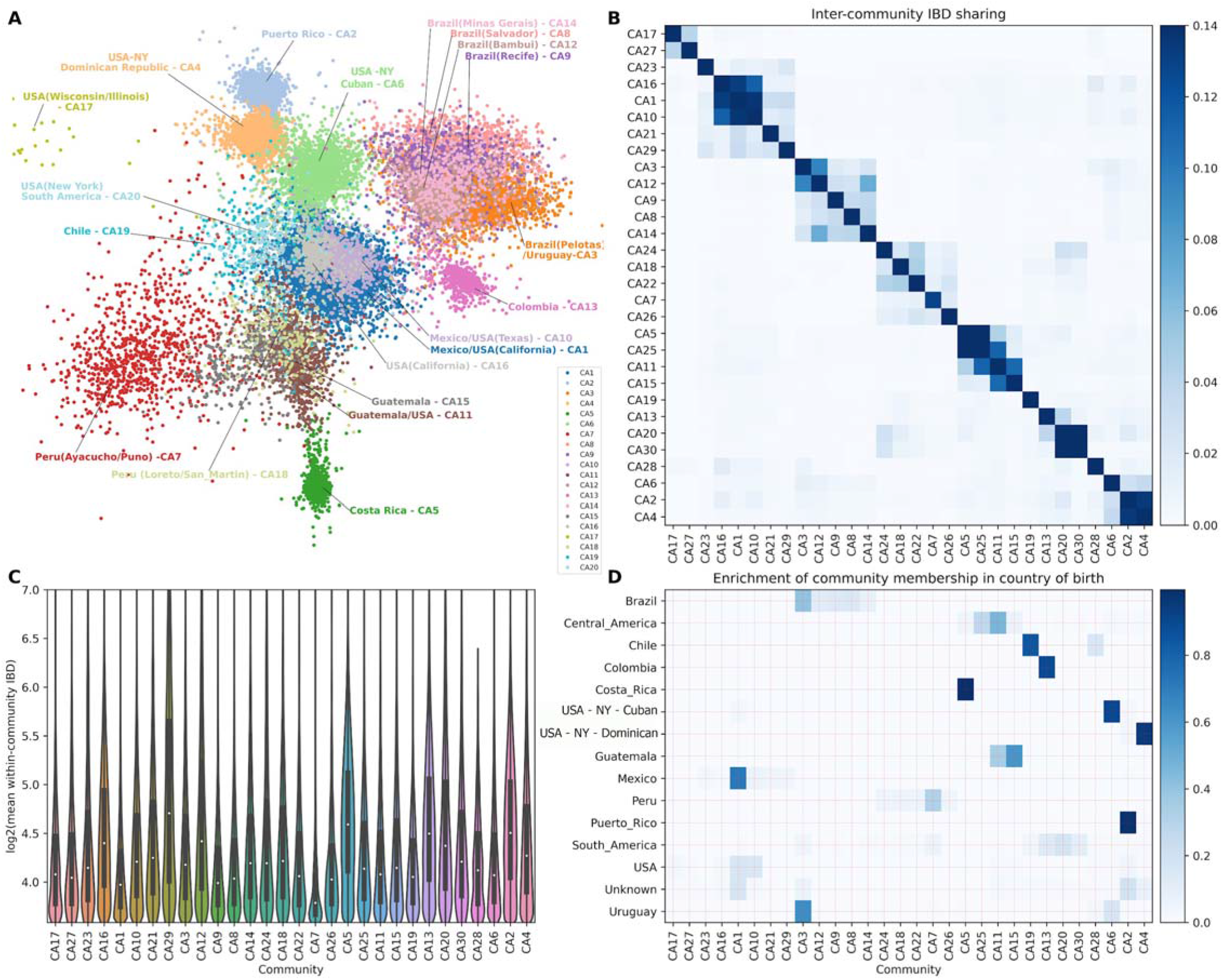
IBD network community detection. We infer the community structure using the infomap algorithm based on a matrix of IBD segments greater than 5cM. A) Top 20 IBD network communities. For visualization purposes, only individuals with connections > 30 are included in the layout calculation. The community labels, such as CA1 and CA2, are named according to the IBD version used and the rank of the community sizes, with CA1 representing the largest community when using all IBD segments. For communities inferred from short and long IBD segments, the corresponding labels are CS1 (Figure S6A) and CL1(Figure S6B), respectively. B) IBD sharing among the top 30 inferred communities (ordered by agglomerative clustering; the same order was followed in C and D). C) Distribution of IBD shared among individuals in each community. D) Enrichment of IBD community membership in the country of origin (i.e., proportions of community labels for individuals born in a given country). To visualize the dynamics before and after the Spanish colonization of the Americas, two different IBD networks were built based on IBD segments between 5-9.3cM (**Figure S6A**) and those > 9.3cM (**Figure S6B**), respectively, which revealed distinct patterns of detected communities.

### Long-distance relatedness among Latin American groups

To explore recent migration among 45 LAm regions, we restricted our analyses to IBD segments greater than 21 cM, representing a recent common ancestor in the last seven generations corresponding to post-colonial times ^45^ and after the admixture process. We reasoned that sharing of larger IBD segments could be originated predominantly from gene flow among regions. At the inter-regional level, we detected higher levels of sharing between Puerto Rico with New York (Specifically with Puerto Ricans in New York) and Hawaii groups. Another tight sub-network of sharing is observed in Brazil (**Figure 5A**), where the South East region (São Paulo and Minas Gerais states) have major connections with other Brazilian populations. Interestingly, there are IBD-sharing connections between Uruguay, South Brazil, and Colombia. On the Pacific side, two Peruvian regions (Ica and Trujillo) show high values of IBD sharing.

**Figure 5.**
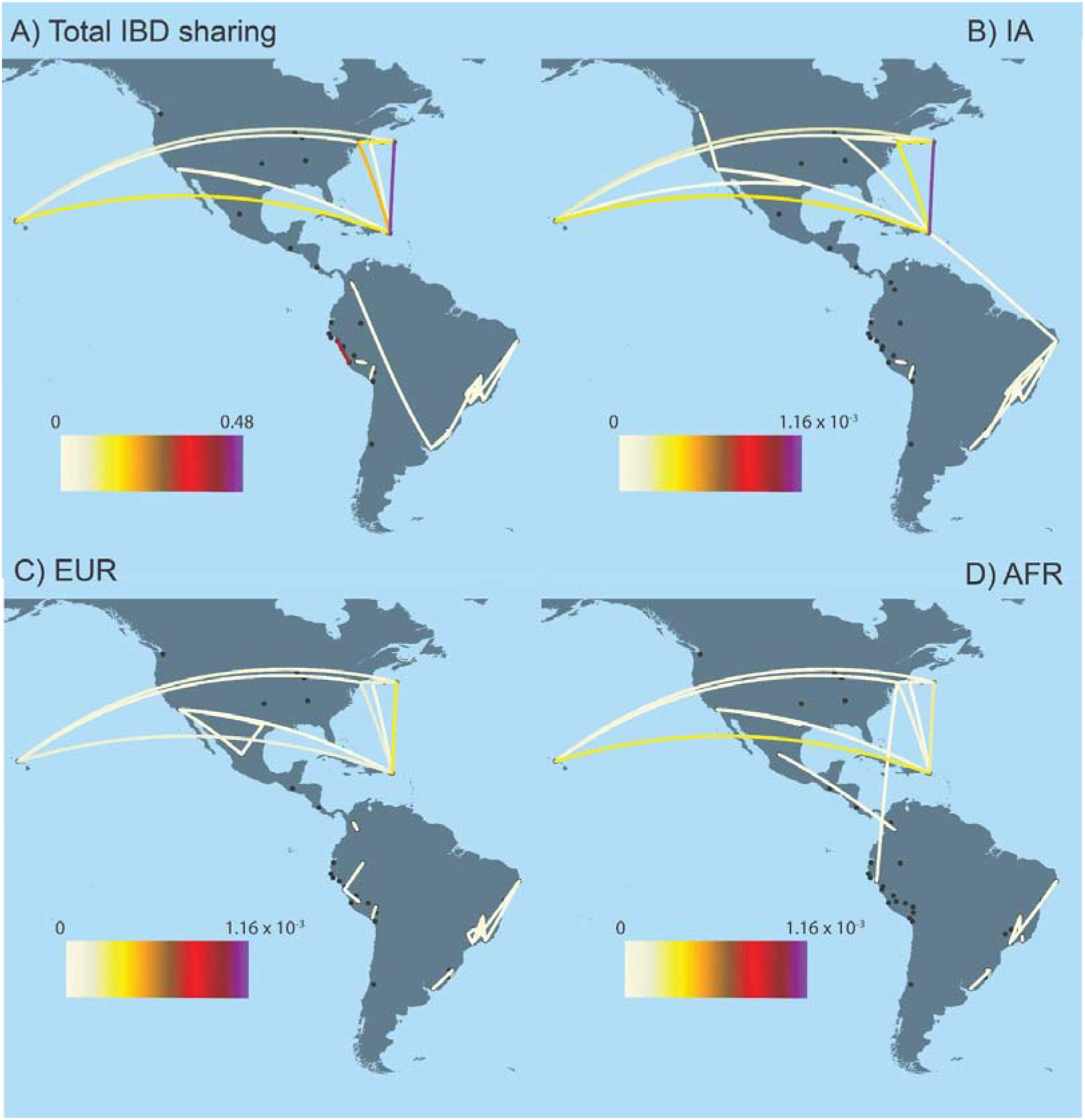
Identical-By-Descent (IBD) analyses of Latin American groups. We explored the relationship among LAm regions by inferring the average IBD shared among regions (A) and an ancestry Specific IBD Score (asIBDScore) for Indigenous American (B), European (C), and African ancestries (D). Dots represent Latin American regions. For African and European Ancestries, we remove the sharing between Peru-Ica and Peru-La Libertad due to their higher sharing and to improve visualization. Plot A range showed the average amount of cM shared among two individuals from populations 1 and 2. On the other hand, the plot range for B-D represents the same statistic focused on segments of a specific ancestry and controlled by global ancestry proportions in each population.

Considering the multi-way admixed origin of LAm populations, we devised a statistic (ancestry-specific IBD score) that quantifies the level of relatedness among two admixed populations for a particular ancestry (AFR, EUR, or IA) (**Figure 5**). We computed the ancestry-specific IBD score (asIBD score, see Methods) by coupling the IBD and local ancestry inferences. Our asIBD score explains the relationship of ancestry-specific IBD segments with respect to the global ancestry of the populations. We detected a different ancestry-sharing pattern between Puerto Rico with New York, and Hawaii (**Table S4A**). A three-way sharing with predominant IA ancestry characterized the sharing among Puerto Rico and New York. On the other hand, the Puerto Rico and Hawaii sharing is characterized by predominant IA and AFR-related ancestries (**Figure 5**). The sharing cluster in Brazil has higher values of asIBD for the IA ancestry, indicating a more homogenous composition of IA ancestry in those regions (**Table S4B**). For IBD sharing between Peru-Ica and Peru-La-Libertad, the EUR ancestry showed the highest value for the asIBD (**Table S4A**).

### Supporting external studies through the GLADdb matching algorithm and statistical genetic software benchmarking

One of GLADdb’s ultimate goals is to provide controls for GWAS and admixture mapping studies. We addressed this goal by developing a genetic matching algorithm. Our method, nearest neighbor simulated annealing matching, shown in **Figure 6A** and outlined in Methods, employs local search to find the optimal cohort from a set of candidates. The algorithm operates on a principal component space in which the external-user-provided query cases can be used to search for controls without needing individual genotypes. The algorithm computes variance-weighted Minkowski distance pairwise between query cases and potential controls, selects the nearest neighbors as candidate controls, samples a set of matches from the candidates, and iteratively resamples and refines the set of matches using simulated annealing, optimizing for the genomic control statistics *λ* ^46,47^.

**Figure 6.**
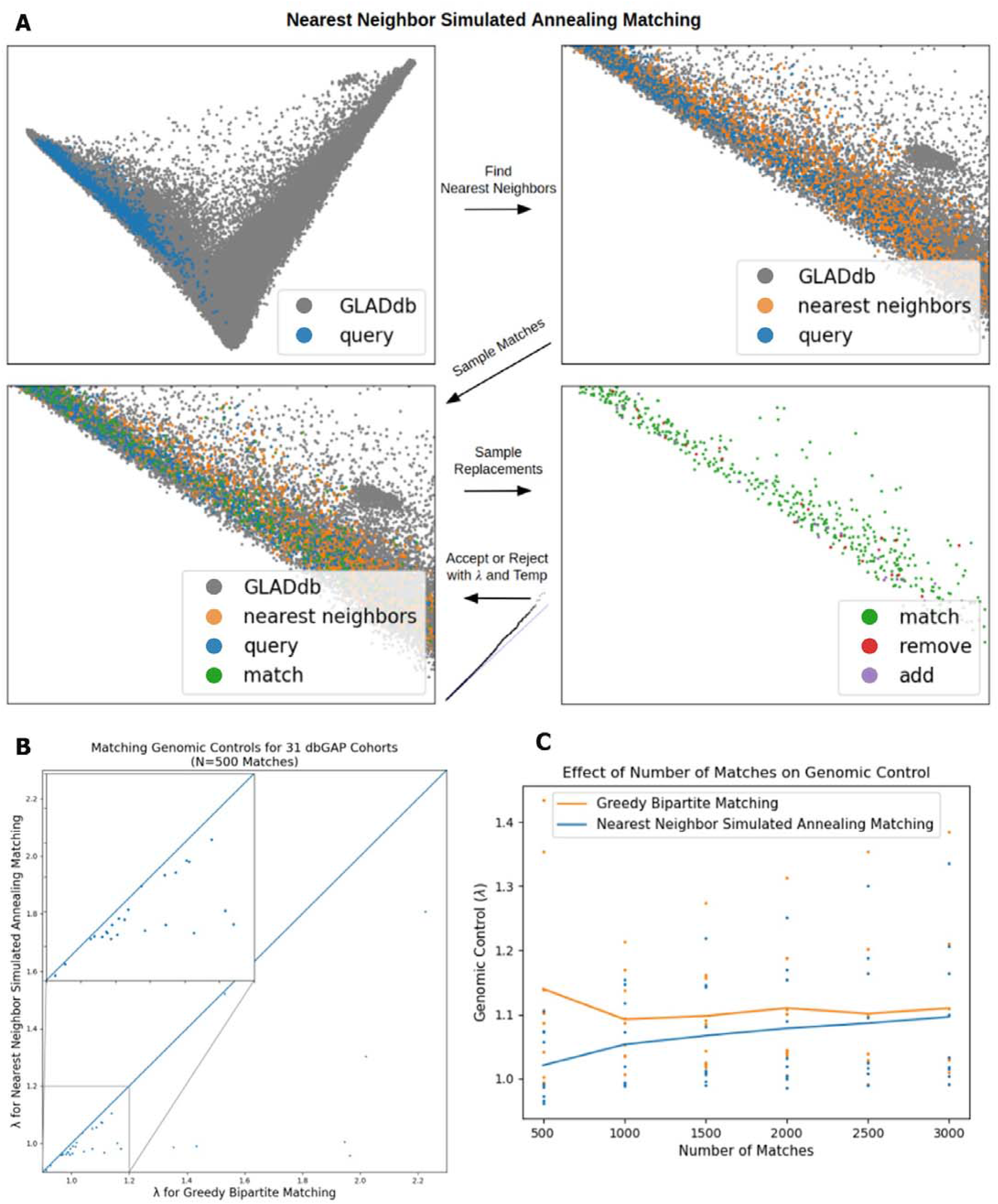
Nearest Neighbor Simulated Annealing Matching Algorithm and Results. A) Visual overview of the algorithm. B) Comparison with baseline bipartite matching algorithm (x-axis), where points below the line y=x indicate our algorithm outperforming the baseline (small box highlights high density region). C) Effect of number of matches on improvement over the baseline.

To evaluate both our matching algorithm and the extent to which GLAD cohorts can provide valid control sets, we performed the following experiment. Using 1000 Genomes populations and some GLAD cohorts as cases, in which the pseudo-phenotype belongs to the query cohort, we ran a greedy bipartite matching baseline ^48,49^ and our matching algorithm and returned summary statistics (i.e., alternative allele frequency, genotype counts, and haplotype ancestry counts by segment) for various control set sizes. Then, for each pair of cases and controls, we ran a GWAS for which the genomic control *λ* statistics are reported in **Table 1** and more extensively in **Figure 6B**. For the analyzed cohorts, which represent a variety of admixed groups, the matched controls yield genomic controls close to 1, suggesting that GLAD will be able to provide useful controls for a variety of cohorts, and our matching algorithm shows slight improvements for larger and more varied query cohorts. These improvements narrow progressively as the number of matches required increases (**Figure 6C**).

**Table 1.** Comparison of genomic control results (*λ* statistics) when returning 500 control individuals from GLAD using the Greedy Bipartite Matching and the Simulated Annealing Nearest Neighbor Matching algorithm.

| Source |  | 1000 Genomes populations |  |  |  | GLAD Cohorts |  |  |  |
| --- | --- | --- | --- | --- | --- | --- | --- | --- | --- |
| Population or Cohort |  | MXL | CLM | PEL | PUR | HCHS SOL | MESA | SIGMA | LARGE-PD |
| Case N |  | 64 | 94 | 85 | 104 | 6646 | 1046 | 1146 | 1463 |
| Greedy Bipartite Matching | Genomic control | 0.9438 ± 0.0009 | 1.0951 ± 0.0018 | 0.9555 ± 0.0012 | 0.9928 ± 0.0017 | 0.9939 ± 0.0063 | 1.0080 ± 0.0136 | 1.0721 ± 0.0074 | 1.0268 ± 0.0177 |
| Nearest Neighbor Simulated Annealing Matching Algorithm |  | 0.9400 ± 0.0001 | 1.0905 ± 0.0009 | 0.9496 ± 0.0010 | 0.9881 ± 0.0003 | 0.9612 ± 0.0045 | 0.9820 ± 0.0047 | 1.0480 ± 0.0072 | 1.0045 ± 0.0050 |

In addition to control matching, GLADdb is an optimal resource for benchmarking statistical genetic software in complex, heterogeneous cohorts with a wide range of available traits. We demonstrated this potential by comparing several popular PRS algorithms (Clumping + Thresholding using PRsice-2^50^, PRS-CS^51^, and PRS-CSx^52^) using a subset of GLAD-SD (**Table S5**, see Methods) with type 2 diabetes (T2D) status, height, or BMI data under a hypothetical scenario where LAm GWAS data is not available (**Table S6**). The GLAD-SD subset includes LAm cohorts with very different population histories and ancestry proportions (e.g., Afro-Caribbeans, Brazilians, and Peruvians). Though the use of the Bayesian PRS-CS method, in general, outperformed PRsice-2, the inclusion of non-European GWAS data using PRS-CSx yielded the largest increase in PRS predictive performance (Figure 7A-C, Figure S7). PRS-CSx improved single-ancestry PRS predictive performance (e.g., East Asian PRS from PRS-CSx versus PRS-CS or PRSice-2) in nearly every instance (**Table S7**). Combining the posterior effect sizes estimated by PRS-CSx further improved models (Figure 7A-C, **Table S7**). Note that the best approach for combining PRS information varied by cohort, likely reflecting cohort heterogeneity (Figure S8). Model performance, as measured by partial R^2^, was negatively associated with mean African ancestry (-0.02 per standard deviation African ancestry, p-value 0.005, Figure 7D). While the percent improvement achieved when leveraging non-European GWAS data can be as high as 80% over the clumping + thresholding model, the R^2^ of each PRS still can be modest. For example, in the Alzheimer’s cohort from the Caribbean, the T2D PRS-CSx model improved prediction by nearly 80%, but the R^2^ of that model was only 0.03 on the observed scale (Figure 7D).

**Figure 7.**
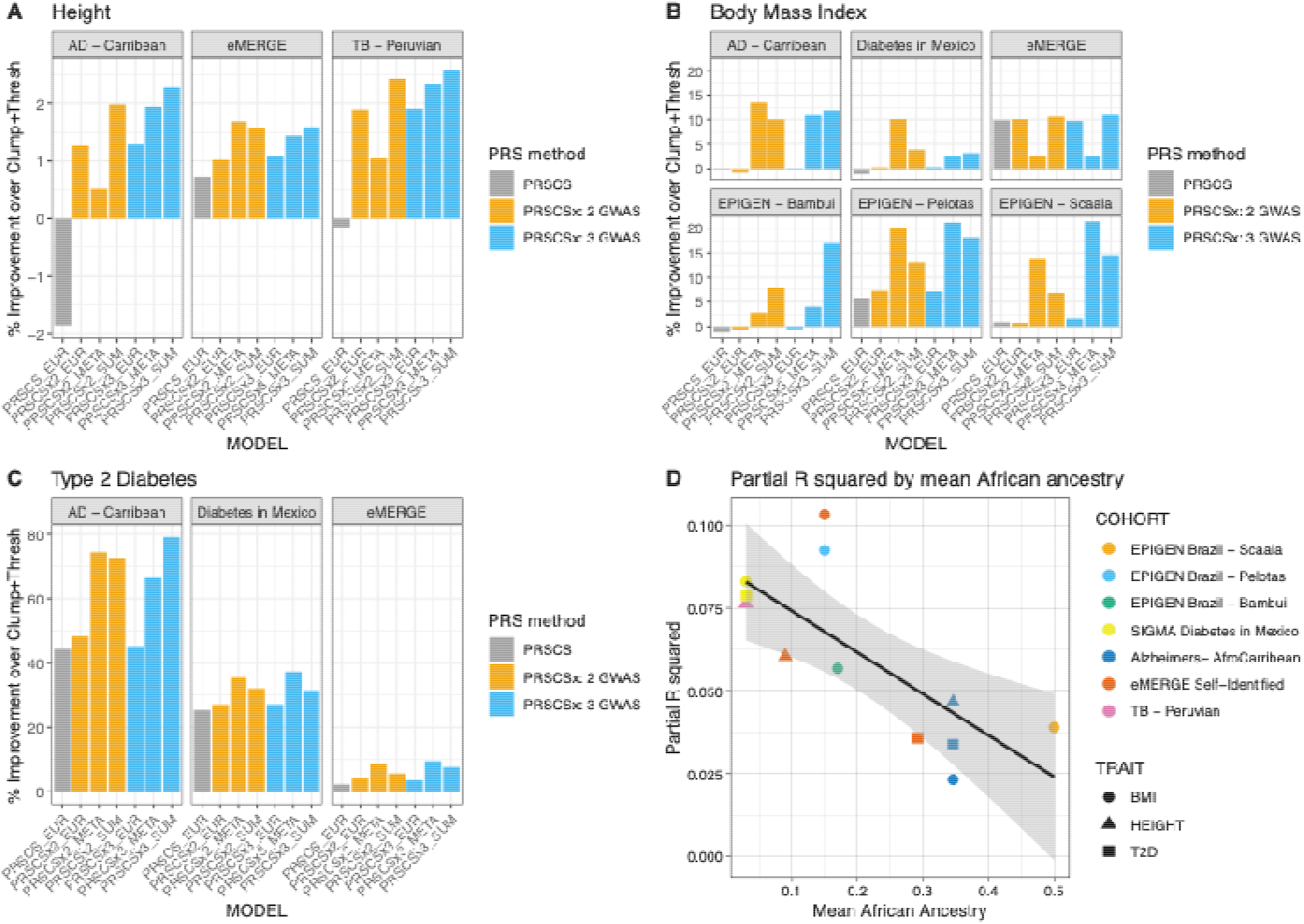
PRS in select cohorts from GLAD-SD. A) Comparison of Height model performance as percent improvement over a European-ancestry GWAS Clumping + Thresholding PRS. Models include PRS-CS using European-ancestry GWAS, PRS-CSx using European and East Asian-ancestry GWAS, and PRS-CSx using European, East Asian, and African-ancestry GWAS. All models were compared using the correlation between the prediction and the trait. B) Comparison of BMI model performance. C) Comparison of T2D model performance. D) Total R2 of best PRS model by African ancestry. Cohorts are labeled by color, traits are labeled by shape. Partial R^2^ was calculated by squaring Pearson’s r followed by subtracting the full model (PRS + covariates) from the base model (covariates only, see methods). African ancestry proportions were estimated using ADMIXTURE.

## Discussion

Latin American individuals are not well represented in genomic and epidemiological studies. This means we have a poor knowledge of their genetic diversity and environmental backgrounds, which limits the applicability of personalized medicine and our understanding of the basis of complex phenotypes ^53^. GLADdb aims to tackle the underrepresentation of genomic data by gathering genome-wide data of LAm populations into a single resource. Through GLADdb, we have two main contributions to LAm genomics: 1) **Population genetics**: we elucidated population structure and gene flow across LAm regions. 2) **Genetic epidemiology**: we developed an algorithm and an online portal (see **Supplementary information 1**) to provide summary statistics from control individuals from GLADdb with a similar genetic makeup to external samples. Also, by assembling a collection of LAm cohorts with very different population histories, we have created a unique tool for evaluating the performance of statistical genetic software in the presence of admixture and other complexities.

For population genetics, continental migrations were the initial sources of LAm diversity. However, other processes have shaped this diversity and the relationships across geographic regions. Through ROH and IBD inferences, we have explored the relationships at intra- and inter-population levels in Latin America in terms of diversity and relatedness. From both analyses, we observed that Peruvians, even with a higher level of homozygosity, have differentiated groups associated with geographical regions ^6,7^. Moreover, IBD sharing tells us more about recent migrations when we restrict the analysis to 21 cM or greater, an interval size correlated with post-colonial events corresponding to the last seven generations before the present. We detected two main networks of sharing: Puerto Rico - New York and Hawaii, and the Brazilian internal sharing groups. In Latin America, during the 20th century, migrations have followed a rural-to-urban or outside-the-country tendency due to regional socioeconomic disparities^54^. Particularly, in Puerto Rico, during the early 1900s, a migration policy was enacted in response to its social and economic problems^55^. Hawaii, Dominican Republic, and Cuba were the primary destinations during the first stage of the Puerto Rican diaspora, followed by a strong migration to New York during the late 1940s ^56^. It is noteworthy that there were socioeconomic differences between the groups participating in each migration stage ^57,58^. For example, many individuals who migrated from Puerto Rico to Hawaii were recognized as *jibaros* ^58^, which are countryside people who farm the land in a traditional way. However, Puerto Ricans who migrated to New York represented a cross-section of economic and social classes ^57^. By inferring the ancestral background of IBD segments, we found that the Puerto Rico/Hawaii sharing is characterized by predominant AFR and IA sharing compared to the IA and EUR sharing between Puerto Rico and New York. These contrasting patterns may reflect the differential composition of the two stages of migration. Brazil is another example of recent migration due to economic factors. During the 1950s, South Eastern Brazil, represented by Rio de Janeiro, São Paulo, and Minas Gerais, experienced a huge economic growth that triggered a massive migration to these regions ^59^. We observed huge connectivity among South Eastern regions (Rio de Janeiro and São Paulo), showing higher values for EUR sharing suggesting higher mobility of European components in Brazil. Moreover, EUR sharing was detected between Southern Brazilian regions and Uruguay. This could reflect their recent shared history as Uruguay was annexed to Brazil before its independence ^60^, and its demographic composition included a significant proportion of Brazilians at that time^61^.

For genetic epidemiology, our genotype matching algorithm and subsequent provision of control summary statistics meet a real need in the research community. Groups exploring the genetic architecture of traits in Latin American cohorts can increase their sample sizes without further straining budgets. This will help facilitate the discovery of genetic risk factors in a historically underrepresented population, which could lead to the discovery of population-specific variation and reduce bias in GWAS data. While there are initiatives that significantly increase the representation of Latin American subjects in genomics, access to that data remains a concern. In some cases, navigating the bureaucratic maze presents a real barrier, while in other cases, the data is proprietary. By constructing the first version of GLADdb, we have already acquired and aggregated Latin American data from across 39 cohorts. In addition, our matching and data transfer processes only require summary statistics (genotype counts and principal components), thus reducing the exposure of sensitive data. Also, by employing our matching algorithm, we can potentially provide a better set of controls than by simply applying for individual cohorts from dbGaP or other public repositories, nor using allele frequencies from heterogeneously sampled cohorts alone.

In addition to supporting genetic studies through control matching, GLADdb presents a valuable resource for evaluating the performance of genetic epidemiology software for methods development and benchmarking. Such software needs to be evaluated in the presence of admixture in addition to the more homogeneous cohorts. This is particularly evident for PRS estimation, where the impact of long-standing biases in GWAS data is well documented ^36,37,62^. In our test case, we evaluated three popular PRS algorithms: clumping + thresholding implemented in PRSice-2, PRS-CS, and PRS-CSx. We found that PRS-CSx, which can model multiple GWAS populations simultaneously, significantly improved predictive performance over single ancestry methods. This was true despite not using GWAS data from any Latin American cohorts for this example. Variability in model performance likely reflected population heterogeneity across the different cohorts, and model performance was negatively associated with mean African ancestry. The sample sizes of the African-ancestry GWAS cohorts used for this study were smaller by an order of magnitude than the East Asian and European Ancestry GWAS cohorts. It is clear that well-powered, diverse GWAS is critical for equitable PRS performance. In the meantime, methodological innovation is required to improve cross-population portability for GWAS traits lacking adequate representation ^63^. In addition to PRS-CSx, several methods such as LDPred-funct and Polypred include functional data, and TL-Multi utilizes transfer learning ^64–66^. The robustness of existing and new PRS methods to admixture can be evaluated using the heterogeneous cohorts represented in GLADdb.

A major challenge in our study, and for LAm genomics, is the poor representation of Indigenous American ancestries. Currently, the Indigenous American representation in public datasets is restricted to a few populations with higher levels of isolation which could lead to caveats in global and local ancestry inferences.

This is important because several studies show that IA ancestry in an admixed LAm population closely relates to their local Indigenous groups ^6,11,16^. To overcome the problems related to IA ancestry, we used a reference panel of Indigenous Peruvians and Guatemalans. These populations have higher effective population sizes compared to other native groups ^67^, which is helpful for avoiding problems related to higher levels of genetic drift. In this way, we can get around the problem of IA inferences in Brazilians or USA individuals with some level of IA ancestry (i.e., Individuals with ancestry related to tribal nations in which genetic studies have not been allowed). Still, better ethically-aware representation in genomics is preferred. Furthermore, GLADdb allowed us to identify geographical regions better represented (e.g., Brazil, Mexico, and Peru) than others in sample size and genotyping technologies (WGS and array data). Moreover, even in these best-represented regions, there is an unbalance of ethnic diversity (e.g., European ancestry descendants are predominant in these datasets). This reality should motivate the need for urgently including regions like Bolivia or Paraguay as well as at the ethnicity level (i.e., African and Asian ancestries in the Americas).

In conclusion, through GLADdb, we highlighted the heterogeneous ancestry composition across LAm populations and inferred ancestry differences in gene flow events relatedness among LAm regions. Also, by sharing summary statistics, we are contributing to improving global equity in genomic research, specifically in epidemiological research in which GWAS is performed routinely. This is one more step to ensuring that health disparities arising from genetic studies do not become pervasive in admixed and non-European populations.

## Acknowledgements

We would like to thank Evangeline “Eevee” O’Connor for assistance in providing an acronym that is both accurate and contributes to generally uplifting our research. TDO was supported by National Human Genome Research Institute of the National Institutes of Health under Award Number R35HG010692. ETS was supported by FAPEMIG (Fundação de Amparo à Pesquisa do Estado de Minas Gerais) RED 00314-16; Programa Nacional de Genômica e Saúde de Precisão – Genomas Brasil from the Brazilian Ministry of Health (CNPq Process 403502/2020-9); Conselho Nacional de Desenvolvimento Científico e Tecnológico - CNPq. RDH is supported by the National Institutes of Health under Award Number R01 GM142112. DPL was supported by the National Heart, Lung, And Blood Institute of the National Institutes of Health under Award Number T32 HL007698-25. JNF was supported by Research Training in the Epidemiology of Aging funded by the National Institutes on Aging under Award Number T32 AG000262.

Latin American Research Consortium on the Genetics of Parkinson Disease (LARGE-PD) is funded by the National Institutes of Health/National Institute of Neurologic Disorders and Stroke (NIH/NINDS) (R01 NS112499).

The NINDS-sponsored Stroke Genetics Network (SiGN) is funded by the National Institutes of Health/National Institute of Neurologic Disorders and Stroke (R01 NS105150 and R01 NS100178).

Genome Sequencing for the Trans-Omics in Precision Medicine (TOPMed) program was supported by the National Heart, Lung and Blood Institute (NHLBI). Genome Sequencing for "NHLBI TOPMed: The Genetic Epidemiology of Asthma in Costa Rica" (phs000988.v3p1) was performed at Northwest Genomics Center (HHSN268201600032I / 3R37HL066289-13S1). Genome Sequencing for "NHLBI TOPMed: San Antonio Family Heart Study (SAFHS) " (phs001215.v4.p2) was performed at Illumina (3R01HL113323-03S1 and R01HL113322). Genome Sequencing for "NHLBI TOPMed: Women’s Health Initiative (WHI)" (phs001237.v3.p1) was performed at Broad Institute Genomics Platform (HHSN268201500014C). Genome Sequencing for "NHLBI TOPMed: Hispanic Community Health Study - Study of Latinos (HCHS/SOL) " (phs001395.v2.p1) was performed at Baylor College of Medicine Human Genome Sequencing Center (HHSN268201600033I). Genome Sequencing for "NHLBI TOPMed: Multi-Ethnic Study of Atherosclerosis"(MESA) (phs001416.v2.p1) was performed at Broad Institute Genomics Platform (3U54HG003067-13S1). Genome Sequencing for "NHLBI TOPMed: Severe Asthma Research Program (SARP)" (phs001446.v2.p1) was performed at New York Genome Center Genomics (3U54HG003067-13S1). Genome Sequencing for "NHLBI TOPMed: Recipient Epidemiology and Donor Evaluation Study-III Brazil Sickle Cell Disease Cohort (REDS-BSCDC)" (phs001468.v3.p1) was performed at Baylor College of Medicine Human Genome Sequencing Center (HHSN268201600033I / HHSN268201500015C). Genome Sequencing for "NHLBI TOPMed: My Life Our Future (MLOF) Research Repository of Patients with Hemophilia A (Factor VIII Deficiency) or Hemophilia B (Factor IX Deficiency)" (phs001515.v2.p2) was performed at New York Genome Center Genomics (HHSN268201500016C). Genome Sequencing for "NHLBI TOPMed: Boston-Brazil Sickle Cell Disease (SCD) Cohort" (phs001599.v1.p1) was performed at Baylor College of Medicine Human Genome Sequencing Center (HHSN268201600033I, HHSN268201500015C, and HHSN268201600033). Genome Sequencing for "NHLBI TOPMed: Children’s Health Study (CHS) Integrative Genomics and Environmental Research of Asthma (IGERA)" (phs001603.v2.p1) was performed at Northwest Genomics Center (HHSN268201600032I). Genome Sequencing for "NHLBI TOPMed: Children’s Health Study (CHS) Effects of Air Pollution on the Development of Obesity in Children (Meta-AIR)" (phs001604.v2.p1) was performed at Northwest Genomics Center (HHSN268201600032I). Genome Sequencing for "NHLBI TOPMed: NHGRI CCDG: The BioMe Biobank at Mount Sinai" (phs001644.v2.p2) was performed at Baylor College of Medicine Human Genome Sequencing Center (HHSN268201600033I) and the McDonnell Genome Institute (HHSN268201600037I). Genome Sequencing for "NHLBI TOPMed: Lung Tissue Research Consortium (LTRC)" (phs001662.v2.p1) was performed at Broad Institute Genomics Platform (HHSN268201600034I). Genome Sequencing for "NHLBI TOPMed: Childhood Asthma Management Program (CAMP)" (phs0017265.v2.p1) was performed at Northwest Genomics Center (HHSN268201600032I). Core support, including centralized genomic read mapping and genotype calling, along with variant quality metrics and filtering, were provided by the TOPMed Informatics Research Center (3R01HL-117626-02S1; contract HHSN268201800002I). Core support including phenotype harmonization, data management, sample-identity QC, and general program coordination were provided by the TOPMed Data Coordinating Center (R01HL-120393; U01HL-120393; contract HHSN268201800001I). We gratefully acknowledge the studies and participants who provided biological samples and data for TOPMed. The full study-specific acknowledgments are included in Supplementary Information 2.

## Methods

### Data Description, Quality control, and imputation

We have gathered data sets for the GLADdb by combining accessible genomic information from Whole-Genome Sequencing (WGS) and microarray genotyping chip sources. We have requested and received access to 39 dbGaP cohorts. Another important source was the WGS projects in TOPMed ^5^. In total, we have explored over 268K samples in detail to find 70,702 Latin American subjects for this initial set. This search includes 172K from general dbGaP datasets including the eMERGE ^68^, PAGE ^69^, and SIGMA ^9^ projects (**Table S1**). **Figure S1** shows our general workflow. For each non-WGS dataset (**Table S1**), we converted their genome coordinates (liftover) from the original reference (NCBI36/hg18 or GRCh37/hg19) to the genome reference GRCh38/hg38 using picard ^70^. After a first liftover run, we used the strand flip option of PLINK ^71^ on the rejected variants and performed a second liftover run. Furthermore, variants were filtered using PLINK for 5% missingness, a p-value less than 1×10^-6^ on the Hardy Weinberg exact test (HWE), keeping only biallelic autosomal variants with a minimum minor allele frequency (MAF) of 1%. Samples were filtered for 5% missingness and heterozygosity exceeding three times the standard deviation from the mean. Also, a linkage disequilibrium (LD) pruned dataset was created using PLINK’s indep-pairwise algorithm using the parameters 50 10 0.1.

For each data set for which we acquired genomic information and appropriate consent, we evaluated self-described demographic variables such as an ethnic designation of Hispanic/Latino. We included the entire cohorts where the primary study design was focused on Latin American individuals, e.g. SIGMA ^9^. For the remaining datasets, many without demographic information provided via dbGaP, we identified possible Latin American individuals using genetic clustering analysis ^39^.

We merged each of these remaining datasets (the LD pruned data) with a custom panel of 361 individuals to assess genome-wide ancestry proportions for European, African, East Asian, and Indigenous American ancestry. This custom panel included 100 each for European, African, and East Asian from the high coverage 1000 Genomes Project data ^38^ (**Table S3**). In addition, we included 61 unrelated, previously estimated as near 100%, Indigenous American high coverage genomes from the Peruvian Genome Project ^6^. Each data set was combined with this reference sample, then we ran a supervised ADMIXTURE analysis ^39^. These results were then evaluated for admixture proportions and any sample found to have greater than 2% Indigenous American ancestry was extracted and included for additional analyses. These samples were then designated as *admixture-defined*, which will persist in our evaluations of the database as to their utility as matches or exclusion.

After we collected all self-described and admixture-defined individuals in each dataset (non-LD pruned data), we imputed the genotype panel against the TOPMed Imputation server ^40^. The TOPMed imputation panel contains over 90K individuals and was shown to accurately impute Latin Americans ^5^. To date, after we have combined across all studies analyzed, including the non-imputed TOPMed WGS data, we have 63,589 non-duplicate samples. This comprises 9,121,629 variants with an imputation r^2^ > 0.3 across all datasets (i.e. no missing data) and includes 8,626,916 SNPs and 494,713 INDELs.

Importantly, GLADdb includes 30,078 individuals with non-ambiguous geographical information (**Table S2**). This means that we have country-level or, in some cases, state or city-level information like Peru, Brazil, and the USA. For the latter three groups, we did not include individuals without state-level information. A particular case is the Rio Grande do Sul state in South Brazil. Two of three cohorts that were sampled in this state correspond to specific cities (Porto Alegre and Pelotas) and were considered as independent groups. To support the clustering of individuals of different project into groups of similar geographical regions (e.g., USA-Wisconsin, Chile, Brazil-São Paulo), we performed an Fst analysis. We calculated the Fst among individuals sampled by different projects but of the same sample region. No regional cluster showed an Fst value above 0.07 (**Table S2**). Finally, these 30K individuals were organized into 45 different regions (**Table S2**). We used this information for ROH and IBD analyses.

After imputation, for each dataset, we kept only variants with r^2^ > 0.9. Then, we merged all datasets and removed variants with missing information in more than 0.1% of the final dataset using bcftools:

~~~
bcftools filter -e ‘F_MISSING > 0.001’ ${mergedGLAD} -O b -o $QC1
~~~

For normalizing and keeping biallelic SNPs we applied the following command line:

~~~
bcftools norm -m +any -s $QC1 | bcftools view -m2 -M2 -v snps | bcftools sort -O b -o $GLAD
~~~

Our initial freeze of GLADdb consists of 3,248,494 biallelic SNPs (r^2^ > 0.9) and 63,589 individuals (**R0.9 dataset**).

To avoid any phase issues during the merging process, we infer the haplotype phase for the complete GLADdb using SHAPEIT ver4 ^72^ using the TOPMed freeze9 dataset ^5^ (160K individuals) as a reference panel. We ran SHAPEIT with the following parameters

~~~
shapeit4 --input $GLAD --map $map --thread 60 --region chr${chr} --reference $TOPMEDRef --output $Phased_GLAD
--log phased_chr${chr}.log --mcmc-iterations 10b,1p,1b,1p,1b,1p,1b,1p,10m
~~~

### Identical-by-Descent and Relatedness analyses

Phased biallelic **R0.9 dataset** together with HapMap genetic maps (GRCh38) were used as input for inferences of IBD (Identical-By-Descent) segments using hap-ibd ^73^. For hap-ibd, we set the parameters “min-seed=3” and “min-output=3” to reduce the rate of false positiveness; defaults were used for all the other parameters. Given IBD coverage is dramatically increased by the paucity of SNP markers, we defined low SNP density regions as 1-cM windows with the number of SNPs less than 30 and processed all IBD segments overlapping with these regions by splitting them and removing the parts within the low SNP density regions. The processed IBD segments were then used as input for ancestry-specific downstream analysis. For non-ancestry specific analyses, we further merged and flattened the processed IBD segments for each sample-pair when two segments are either overlapping or close (gap no longer than 0.6cM and the number of phasing-informative discordant markers no more than 1) ^74^. The flattened and merged IBD segments were kept if the segment length >= 5cM. Genome-wide total IBD length of all segments shared by each sample pair was then calculated and organized to an IBD matrix with each element representing the relatedness between a pair of individuals. For agglomerative clustering, we transformed the matrix into a dissimilarity matrix by the formula X = (max-min)/(X-min+1e-9). The IBD post-processing steps including encoding, removing low SNP density regions, decoding, sorting, merging, filtering, and matrix-building were implemented in a C++ toolkit *ibdtools* (https://github.com/umb-oconnorgroup/ibdtools) to accelerate the computation for large IBD datasets, for instance, hundreds of billions of IBD segments.

We estimated the kinship coefficient for each pair of individuals in GLADdb with IBDkin ^75^. After kinship coefficient inferences, we pruned for relatedness in GLADdb using NAToRA ^76^ to exclude the minimum number of related individuals while removing the main kinship relationships in the dataset. We used 0.03125 as the kinship coefficient threshold which is the theoretical kinship coefficient expected for a 4th degree relationship.

### Continental Population Structure

#### PCA and UMAP

Prior to performing dimensionality reduction, we used PLINK^71^ to narrow our biallelic **R0.9 dataset** by applying LD pruning with a threshold of 0.5 to all 54K samples. Then, using the scikit-learn ^77^ implementation, we ran Principal Components Analysis (PCA) on the LD-pruned sites, keeping the top 50 components. To help with cluster visualization, we reduced the 50 principal components down to 2 dimensions by applying the UMAP algorithm, using the umap-learn package ^78^, with n_neighbors set to 10 and min_dist set to 0.25.

#### Runs of Homozygosity (ROH)

We inferred the ROH segments for our 45 Latin American groups and 21 reference populations to explore the level of homogenization in each group. For each group, we used PLINK to filter for monomorphic variants and generate a transpose format. Then, we ran GARLIC ^44^, software that infers ROH based on the Pemberton *et al* ^79^ pipeline detecting short (tens of kb), medium (hundreds of kb to several Mb) and long (tens of Mb) ROH segments. We set the --auto-winsize mode to allow GARLIC to estimate the best window size for ROH inference starting from a 50 SNP window. We used the following command line:

~~~
garlic --tped ${pop}.tped --tfam ${pop}.tfam --build hg38 --error 0.001 --cm --winsize 50 --auto-winsize --
auto-winsize-step 10 --out roh_autosize_${pop} --threads 20 --map ${geneticmap}
~~~

For each individual in each group, we summed all ROH sizes to determine the genome-wide amount of ROH. Considering the good representation of Peruvian and Brazilian regions, 13 and 12 respectively, we estimated the correlation between the median for each group and the average genome-wide ancestry proportion in each country using the Pearson correlation. Processing and plotting scripts are available in: https://github.com/umb-oconnorgroup/GLAD_DemographicAnalysis

#### Local ancestry Inferences

We ran local ancestry inference using RFMix ver2 ^80^ on GLADdb. We inferred local ancestry for the phased dataset considering two Expectation-Maximization runs and eight generations since admixture. For the ancestry reference panel, we selected 982 individuals including 250 Europeans, 250 East Asian, 250 Africans and 232 individuals with predominant Indigenous American ancestry (Table S2). Europeans, Africans and East Asian reference populations are part of the 1000 Genomes Project. Individuals with predominant Indigenous American ancestry includes Indigenous Americans from the Peruvian Genome Project ^6,7^ and individuals with predominant Indigenous American ancestry (above 99% of Indigenous American ancestry) from Guatemala (Table S2).

### Distant genetic relatedness among Latin American groups

#### IBD-community detection

For community detection, we calculated an IBD matrix by summing up all IBD segments with length within a specific range (>5cM, 5-9.3 or >9.3cM) across the genome for each pair of individuals, and set all elements with values < 12 cM to 0 in this matrix to reduce the density of non-zero elements in the matrix. The resulting symmetrical matrix was used as a weighted-adjacency matrix to build a bidirectional relatedness network. We used the infomap algorithm implemented with the python-igraph ^81^ package to infer the community structure of the relatedness network. We kept individuals within the top 20 communities and with a degree >= 30 connections and used the Frutcherman Reingold layout ^82^ for visualization purposes. Community enrichment in a given birth country is defined as the largest proportion of community labels for individuals born in the country. The number of communities enriched in a birth country is determined by counting the communities that have >1% enrichment in this country.

#### IBD sharing among Latin American regions

To explore the recent relationship among Latin American regions, we focused on IBD segments greater than 21.4 cM. We calculated the IBD sharing at intra and interregional levels. For intraregional sharing, we summed the total amount of shared IBD and divided it by the number of pairs: N(N-1)/2, where N is the total number of individuals included for that region. For interregional sharing, we summed the total amount of shared IBD among individuals of populations 1 and 2 and divided it by N_1_xN_2_, where N_1_ and N_2_ are the total number of individuals included for populations 1 and 2 involved in the sharing, respectively.

#### Ancestry Specific IBD

From the multi-way admixed origin of Latin American populations, IBD (segments greater than 21.4 cM) and local ancestry analyses provide an opportunity to detect ancestry-specific signatures related to bottleneck (whitin-region analysis) and recent migration (across-region analysis) along the Americas.

We implemented a python algorithm called *GAfIS* (that stands for “Getting Ancestry For IBD Segments”) that uses RFMIX outputs to identify local ancestry labels for an IBD segment shared by a pair of individuals under a certain probability threshold. As a probability threshold for local ancestry inferences in *GAfIS*, we set 90% for a genomic region being of the K ancestry. For this analysis, we included our processed IBD segments to reduce the proportion of false positives. Moreover, if an IBD segment contained several ancestries, we split the segment into segments corresponding to independent ancestries for each pair of individuals.

After ancestry identification of the IBD segments, we filter out ancestry specific-IBD segments based on the following criteria:

-The ancestry profile of one of the individuals for the IBD region was unknown for having a local ancestry probability lower than 90%.

-Both individuals have different ancestry labels of the IBD segment.

After those filters, we kept individuals with demographic information and calculated an *ancestry-specific IBD score* (asIBD score) within and across the 45 Latin American groups. Our asIBD score is defined in the following equations:

Within regions:

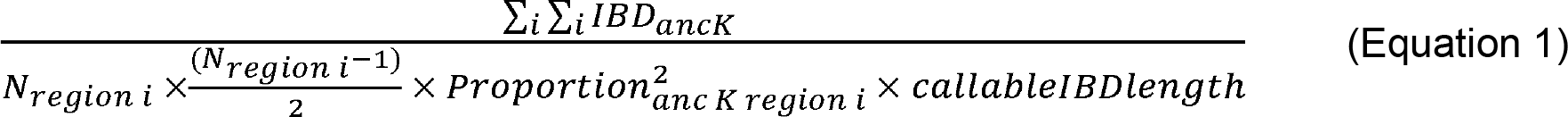

Across regions:

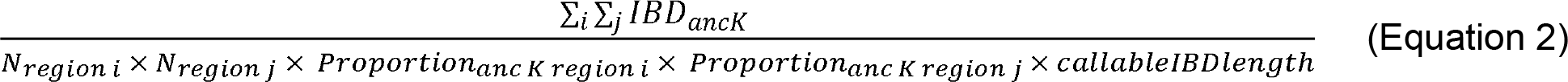

Where:

Anc K= African, European or Indigenous American ancestries

IBD _anc K_: The total amount of ancestry K IBD shared between a pair of individuals from region i and j.

N _region i_: Total number of individuals from region i.

N _region j_: Total number of individuals from region j.

Proportion _anc K region i_: Global ancestry proportion for Ancestry K in region i.

Proportion _anc K region j_: Global ancestry proportion for Ancestry K in region j.

callableIBDlength: Total size of the genome that was included for IBD analysis.

In both equations, in the numerator, for a specific ancestry, we summed the total amount of IBD per ancestry for each pair of individuals from the same region (Equation 1) or between region i and j (Equation 2). To control for sample size and ancestry proportions, for equation 1, we divide the total amount of shared IBD by the product of the total number of combinations of individuals and the square of ancestry proportion. For Equation 2, we divide by the product of sample size for each region and the product of the global ancestry proportion K for each region, respectively. Finally, to get a value relative to the total size of the genome, we included the genome size that was analyzed in the IBD inference in both equations. Codes and pipeline to estimate the asIBD score are available in: https://github.com/umb-oconnorgroup/GLAD_DemographicAnalysis

### Polygenic Risk Scores in Latin American populations

#### Description of PRS cohorts

We utilized the following studies participating in GLAD: Columbia University Study of Caribbean Hispanics and Late Onset Alzheimer’s disease (phs000496), Slim Initiative in Genomic Medicine for the Americas (SIGMA): Diabetes in Mexico Study (phs001388), eMERGE Network Phase III: HRC Imputed Array Data (phs001584), Early Progression to Active Tuberculosis in Peruvians (phs002025), and EPIGEN-Brasil (Bambui, Pelotas, and SCAALA). These studies all ascertained one or more of the following traits: height, body mass index (BMI), and/or type 2 diabetes (T2D). See Table S5 for a complete description of cohorts.

#### Ancestry proportions, relationship inference, principal components and imputation

Within each cohort, PCs were calculated using PC-Air ^83^ to utilize as covariates. Related individuals were resolved to the 3rd degree using a kinship matrix generated in *Identical-by-descent and relatedness analyses* section. Genotyped data from each cohort was separately merged with the 1000 Genomes Project (1KGP) ^38^. Global ancestry proportions were estimated using ADMIXTURE^39^, a K of 5, and 20 replicates. For PRS estimation, imputed variants were filtered for a minimum imputation r^2^ of 0.9 and a MAF of 0.01. Both imputed and genotyped data were down-sampled to Hapmap Phase Three variants as required by PRS-CS ^51^. Phenotype data was harmonized across cohorts, though all analyses were conducted on a per-cohort basis.

#### GWAS summary statistics

Genome-wide association statistics were obtained from the GWAS Catalog ^85^, Biobank Japan ^30^ (BBJ), and UK Biobank ^28^ (UKBB). African-ancestry GWAS summary statistics were combined using a random-effects meta-analysis using the GAP package in R to improve sample size. See Table S6 for a description of summary statistics used for this study.

#### Heritability estimation

Per-cohort additive heritability for each trait was estimated using GCTA ^86^, adjusting for sex, age, age^2^, and PCs 1-10. For each set of GWAS summary statistics, heritability was estimated using LD score regression ^87^, using the appropriate 1KGP super-population for the calculation of LD scores.

#### Polygenic risk score calculation

Pruning/Thresholding PRS: We used PRS calculated with PRSice-2 ^50^ as the representative pruning and thresholding (P+T) method. For P+T, we trained the r^2^ parameters (r^2^ thresholds of 0.2, 0.4, 0.6, and 0.8), window size (+/- 250 kb, 500kb, 750kb, 1000 kb), and p-value thresholds (iterated by PRSice-2) in one cohort (eMERGE) and validated the parameters in the other cohorts.

Bayesian Mixture PRS: We used PRS estimated with PRS-CS^51^ as the baseline Bayesian mixture method. For PRS-CS, we trained the phi (φ) parameter (phi=1e-06, 1e-04, 1-e02, and 1e+00) in one cohort (eMERGE, as this cohort included information for all tested traits) via a small grid search and validated it in the other cohorts. In addition, we also evaluated the fully Bayesian pseudo-validation method (phi=auto) for obtaining phi.

Multi-ancestry PRS using PRS-CSx: We leveraged PRS-CSx ^52^ to compute a multi-ancestry PRS, which simultaneously fits multiple sets of GWAS summary statistics while modeling population-specific LD, resulting in more accurate posterior effect sizes for any relatively underpowered GWAS. PRS-CSx outputs a PRS corresponding to each GWAS population and an inverse variance meta-analysis of the posterior effect sizes. We trained the best linear combination of each single-population PRS in one cohort using the mixing weights method proposed by Márquez-Luna et al. ^88,89^ (Equations 3 and 4) with validation in other cohorts. Prior to combining, each PRS is scaled (mean 0, standard deviation 1). In addition, we also evaluated weighting PRS by ancestry proportions (Equation 5), weighting by ancestry proportions after collapsing East Asian and Indigenous American ancestries (Equation 6), and regressing on ancestry proportions prior to model fitting. We compared these linear combinations to the PRS generated from the inverse-variance meta-analysis of PRS-CSx posterior effect sizes.

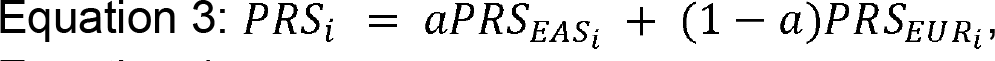

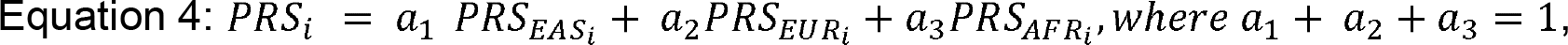

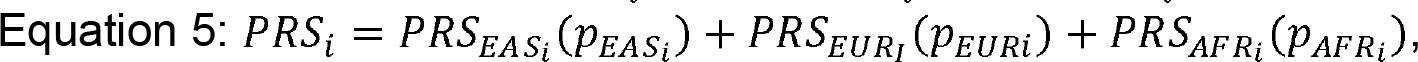

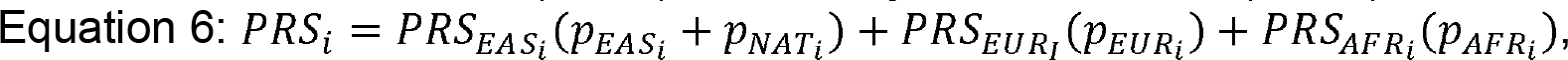

where *α*, *α_1_*, *α_2_*, and *α_3_* represent mixing weights, *PRS_AFR_i__*, *PRS_EUR_I__*, and *PRS_EAS_i__* represent a PRS calculated using African, European, East Asian ancestry GWAS, respectively, for individual *i*. *p_EAS_i__*, *p_AFR_i__*, *p_EUR_i__*, and *p_NAT_i__* represent the East Asian, African, European, and Indigenous American ancestry proportions for individual *i*.

For BMI, height, and T2D, GWAS summary statistics from East Asian, European, and African populations are publicly available (see Table S6). In addition, we were able to train the full range of parameters thanks to multiple independent Latin American cohorts containing data for these traits. We first compared pruning and thresholding (P+T), PRS-CS, and PRS-CSx models. We then evaluated PRS-CSx based multi-ancestry models, comparing linear combinations (the best performing linear combination model for each cohort) and inverse-variance meta-analyses of PRS-CSx posterior effects. These multi-ancestry models were derived from East Asian and European GWAS (referred to as SUM2 and META2) or derived from East Asian, European, and African GWAS (referred to as SUM3 and META3). Finally, we compared these multi-ancestry models against the best single ancestry PRS (EUR2 and EUR3 estimated using PRS-CSx).

#### PRS model evaluation

All models were evaluated using the 10-fold cross validation framework outlined by Pain et al ^90^. In this approach, the primary metric is the Pearson correlation between the predicted and true values with a standard error of 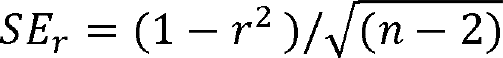, where r is the Pearson correlation and n is the sample size. Correlations were compared using the two-sided William’s test implemented in the psych R package that accounts for the non-independence of the model predictions. R^2^ was calculated as the square of Pearson’s r; partial R^2^ was estimated by subtracting the R^2^ of the base model (only covariates) from R^2^ of the full model (covariates and PRS). In general, the base model included age, age^2^, sex, and PCs 1-10 with the exception of cohorts with a categorical age variable (eMERGE for T2D). The Pelotas cohort, as a birth-year cohort, age and age^2^ were not included as all subjects were the same age. We tested the association of mean ancestry proportions of the cohorts with model performance using linear regression, adjusting for the GWAS trait (R2∼ scaled ancestry proportion + trait).

#### Code Availability

From the code utilized for this project, we developed an R package called PRSHelpDesk that supports PRS estimation and evaluation. It is available on GitHub at https://github.com/dloesch/PRSHelpDesk.

### Matching

Both the baseline bipartite matching ^49^ algorithm and the nearest neighbor simulated annealing matching algorithm operate on a principal components space composed of the first 50 components computed using 246,799 LD-pruned SNPs from GLADdb. The external-user-provided query is also embedded into the PCA space with a saved transformation matrix and pairwise distances are computed with a variance-weighted Minkowski distance metric. Once a suitable matching set has been found, we return summary statistics to the external user including alternate allele frequency, genotype counts, and haplotype ancestry counts by segment.

The baseline algorithm is outlined in **Algorithm 1** and consists of iteratively applying scikit-learn’s ^77^ bipartite matching implementation until enough controls have been found.

Given a desired control cohort size *m* and hyperparameters *α*, *β*, *γ*, and *n*, the nearest neighbor simulated annealing matching algorithm, outlined in **Algorithm 2**, proceeds as follows. The computed pairwise distances between query and GLADdb PCA embeddings are used to find the *α* nearest neighbors of each query genome from the potential controls, which we then merge into a candidate set. We sample *m* controls from the candidate set and do so *β* times to generate *β* control cohorts. We use the genomic control *λ*, calculated between a control cohort and the query, to evaluate the *β* control cohorts. The *λ* values are then used to select the optimal starting control cohort and a function of their standard deviation is used to initialize our simulated annealing temperature. We perform simulated annealing for *n* iterations, randomly swapping *γ* genomes between our control cohort and the candidate set at each iteration, evaluating the control cohort by its genomic control *λ*.

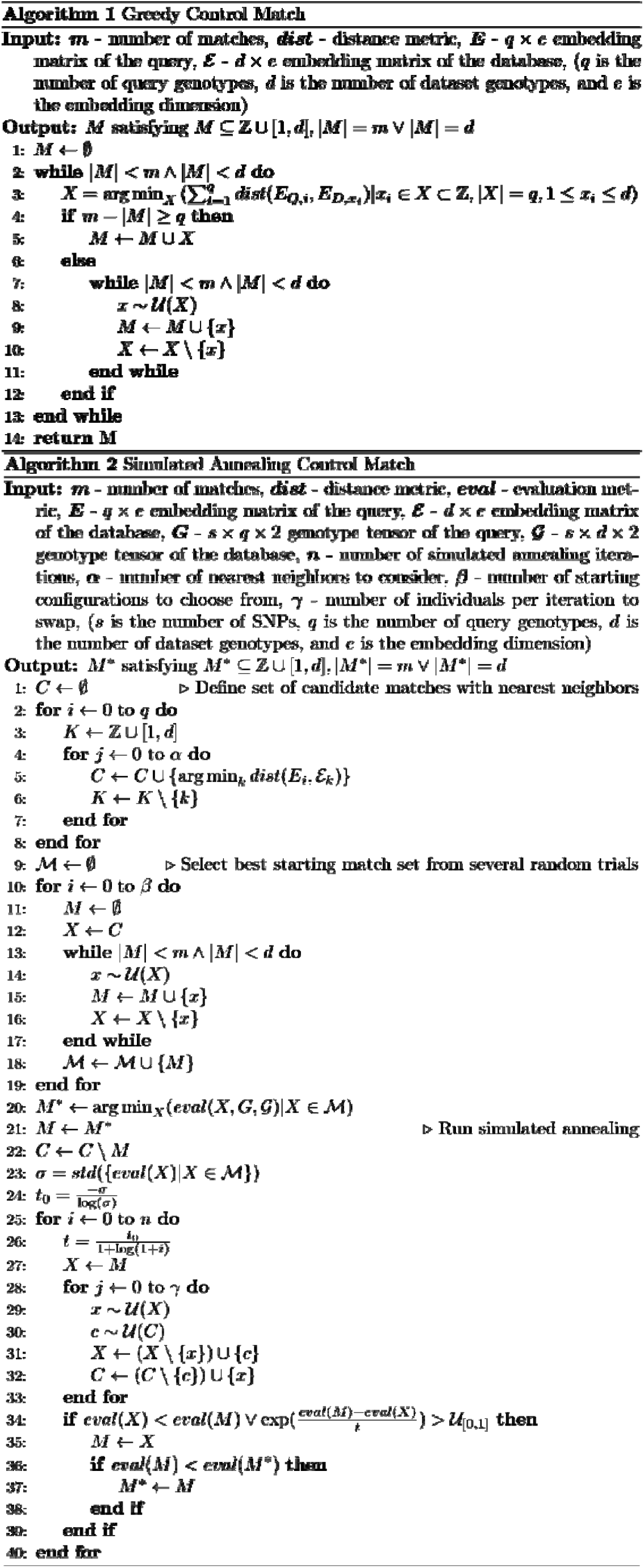

### GLADdb

The GLADdb portal provides both visualization and control matching functionality. The visualizations are built with the Plotly library, enabling in-browser interaction, zooming, and filtering. The control matching page enables filtering by self-identified ethnicity, phs numbers, and some phenotypic traits. The external user is asked to prepare and anonymize their data using a Dockerfile provided at github.com/umb-oconnorgroup/gladprep.

### Code availability

The code utilized for this study is publicly available on GitHub at:

IBD analysis: https://github.com/umb-oconnorgroup/ibdtools

ROH and Ancestry Specific IBD: https://github.com/umb-oconnorgroup/GLAD_DemographicAnalysis

PRS Estimation and Evaluation: https://github.com/dloesch/PRSHelpDesk.

GLADdb: https://github.com/umb-oconnorgroup/gladprep and https://github.com/umb-oconnorgroup/gladdb

### Data availability

No data were generated for this study

## SUPPLEMENTARY FIGURES

**Figure S1.**
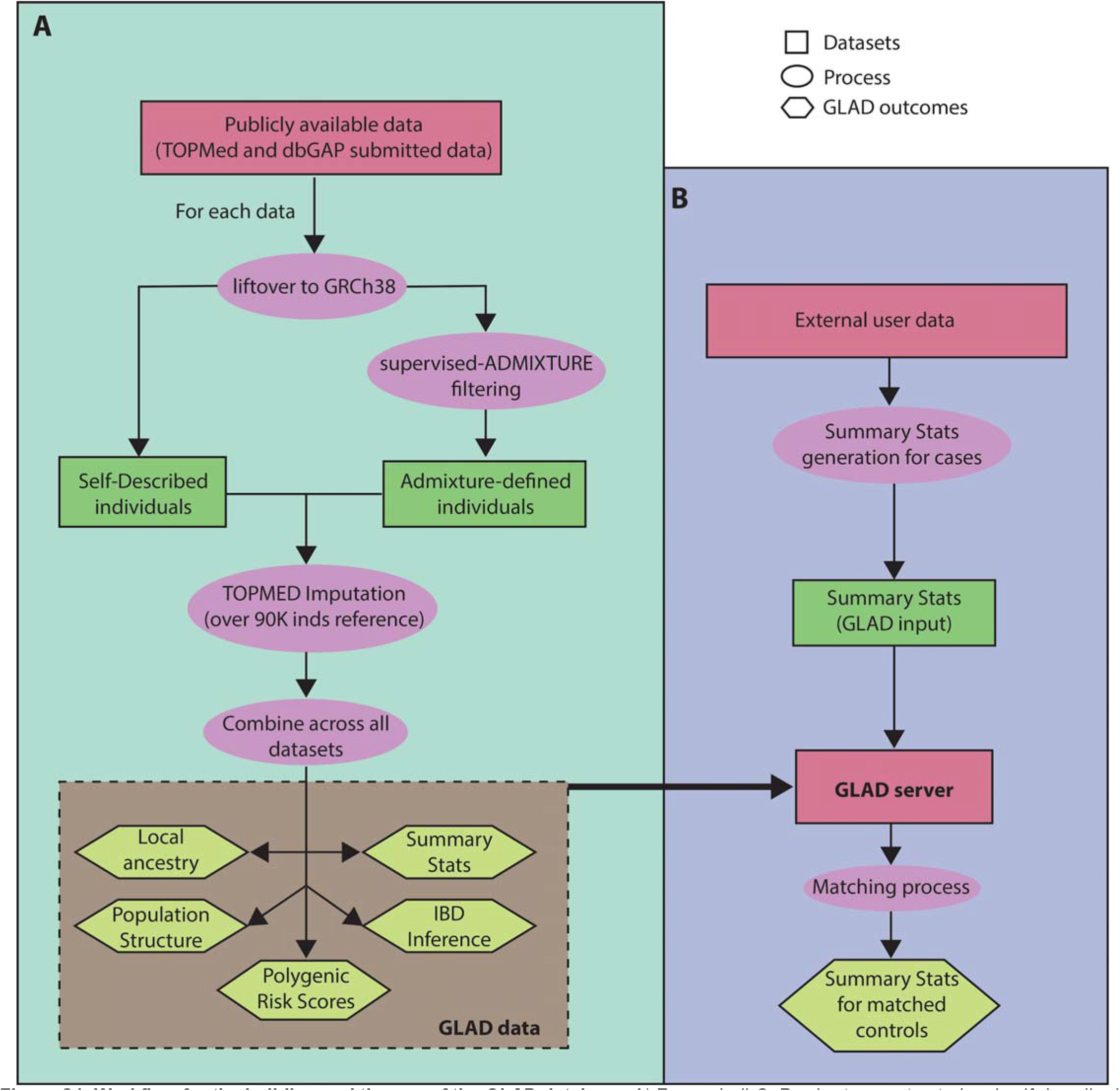
Workflow for the building and the use of the GLAD database. A) For each dbGaP cohort, we extracted and self-described Latino and ADMIXTURE defined subjects with at least 2% of Indigenous American ancestry. Then each cohort was imputed in the Michigan Imputation Center using the TOPMED Imputation panel. After imputation, we selected the best imputed loci (r^2^>0.9) and merged the data. We characterized the GLADdb using PCA, IBD and local ancestry analyses. B) By identifying the GLAD individuals that have similar genetic patterns of a query sample, we provide summary statistics of the control subjects from GLADdb.

**Figure S2.**
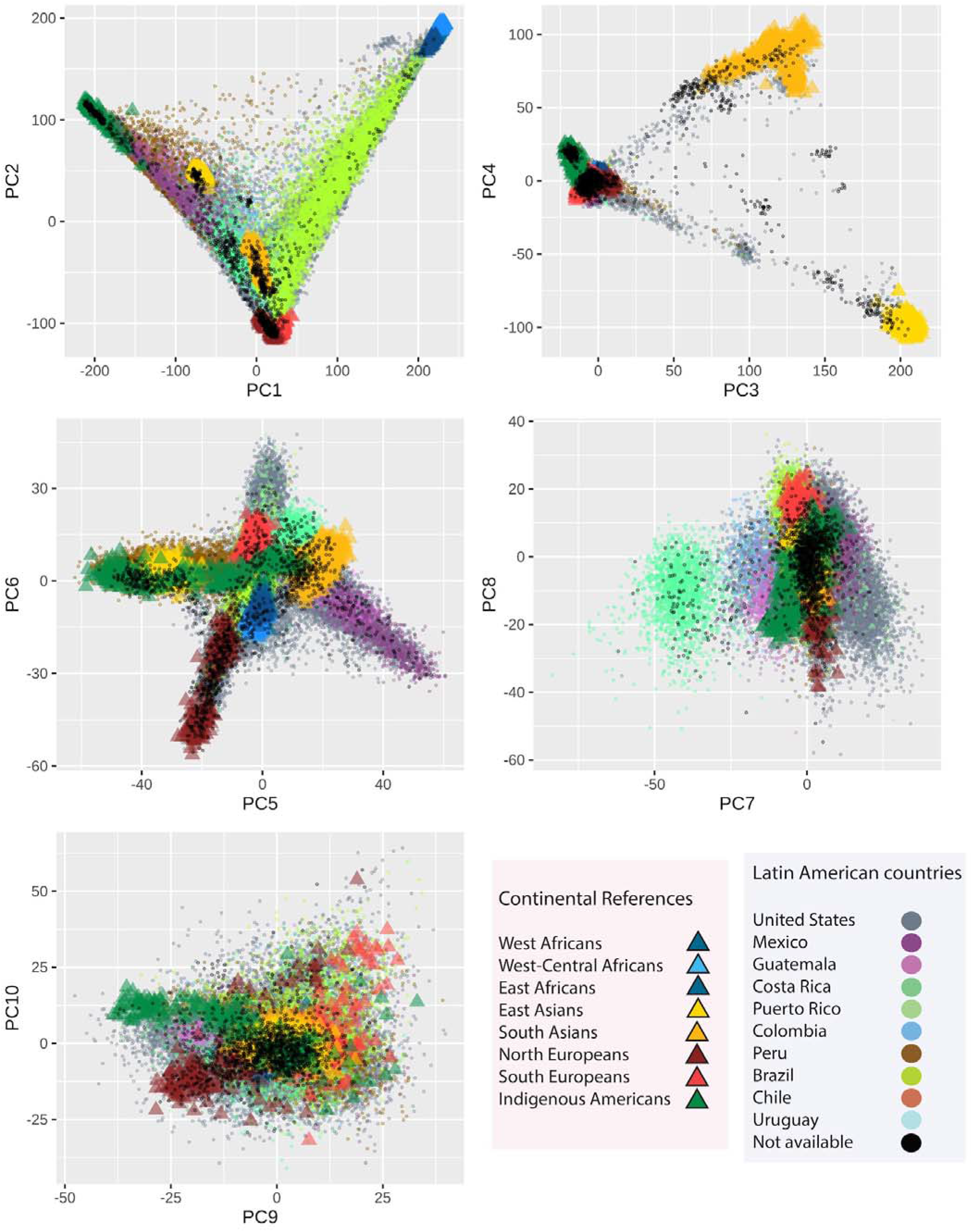
Principal component (PC) analysis GLADdb and ancestral reference groups individuals. First ten PCs that include reference groups (triangles) and GLAD individuals (circles).

**Figure S3.**
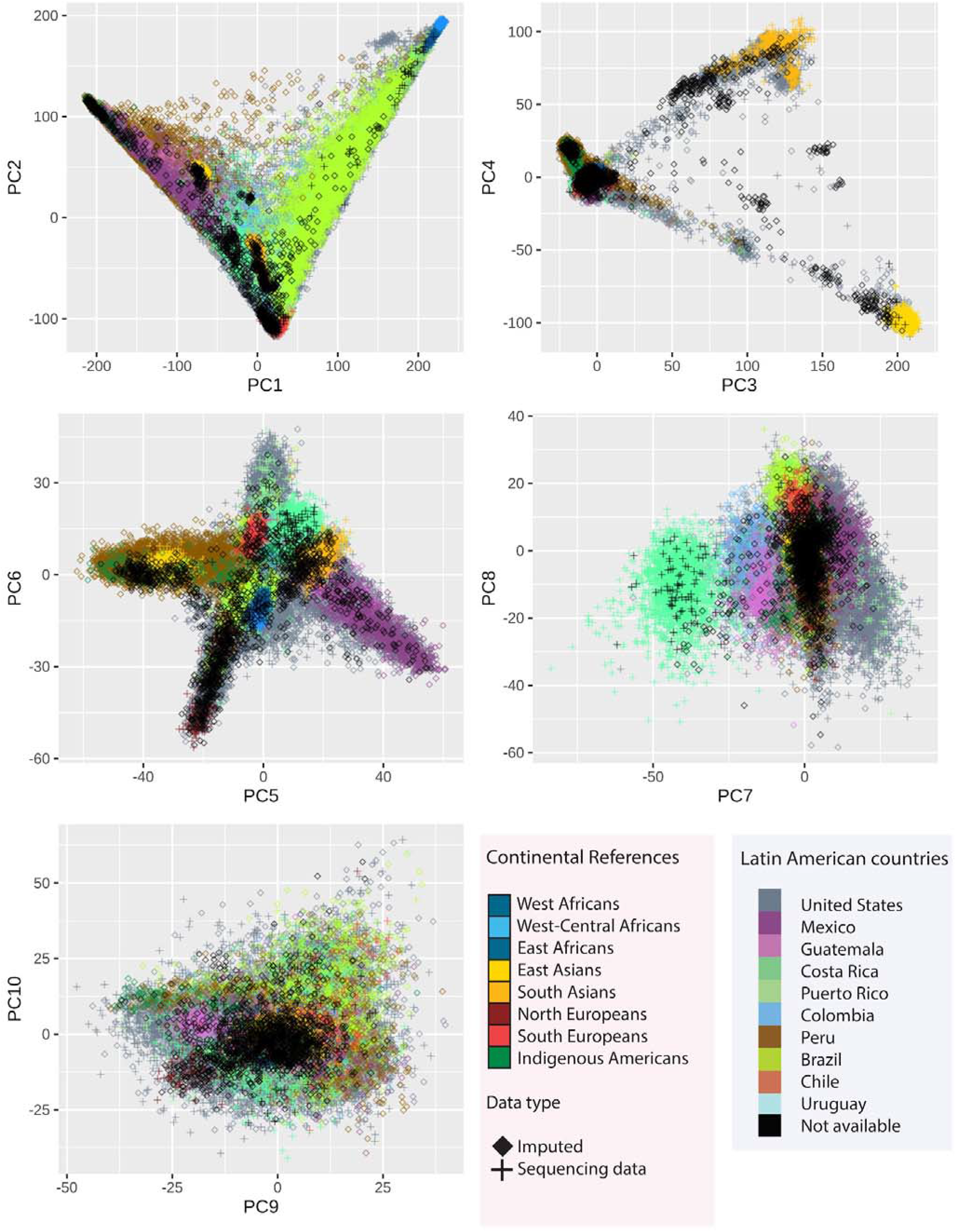
Principal component (PC) analysis GLADdb and ancestral reference groups individuals. Plot shows the relationships between GLADdb individuals with different data types: Imputed (diamond) and sequencing data (cross).

**Figure S4.**
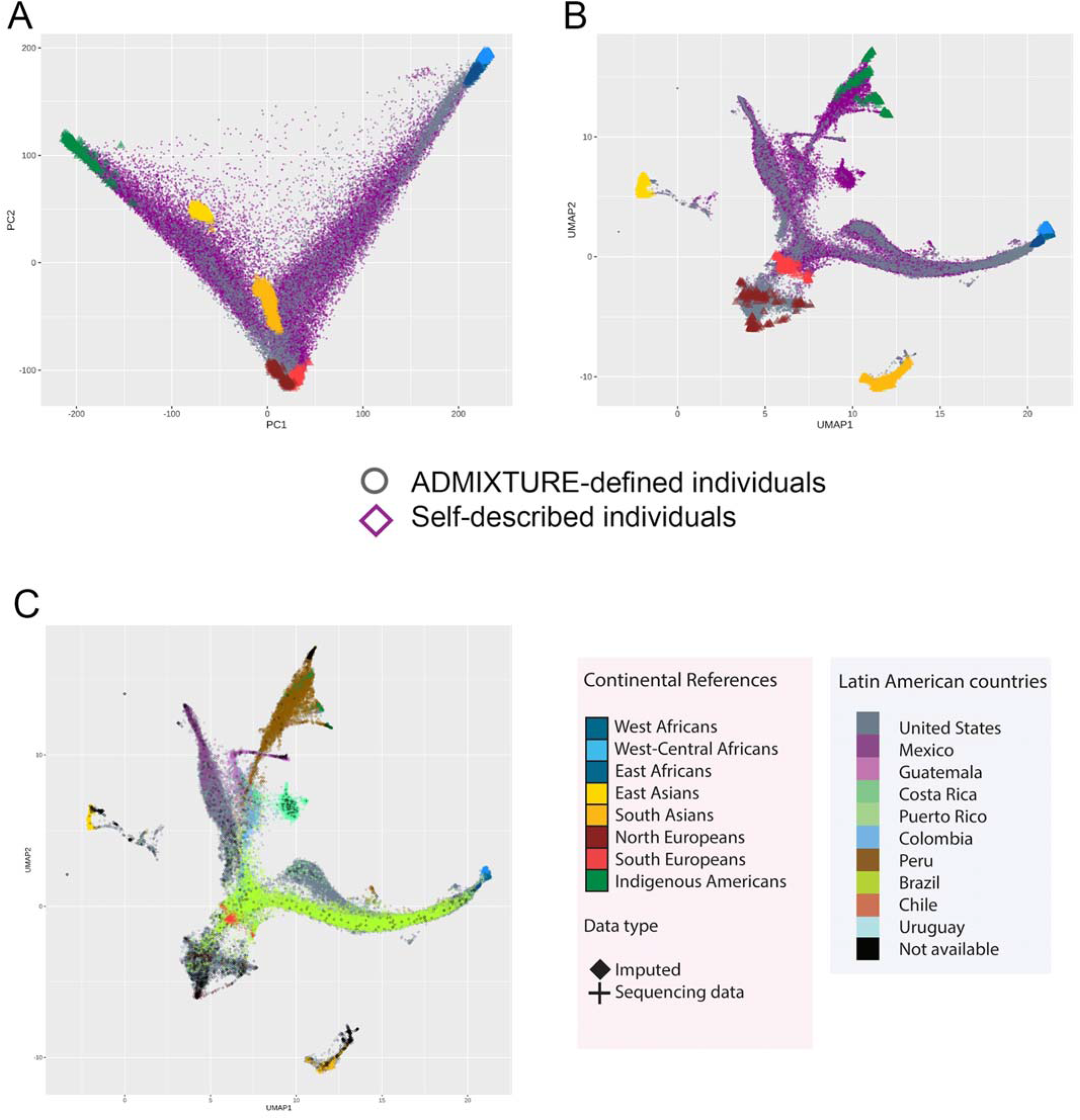
Population structure analysis GLADdb and ancestral reference groups individuals. Principal component (A) and UMAP (B) analyses showing the relationship between self described and ADMIXTURE-defined individuals in GLADdb. C) UMAP analysis of GLADdb individuals with different data types: Imputed (diamond) and sequencing data (cross).

**Figure S5.**
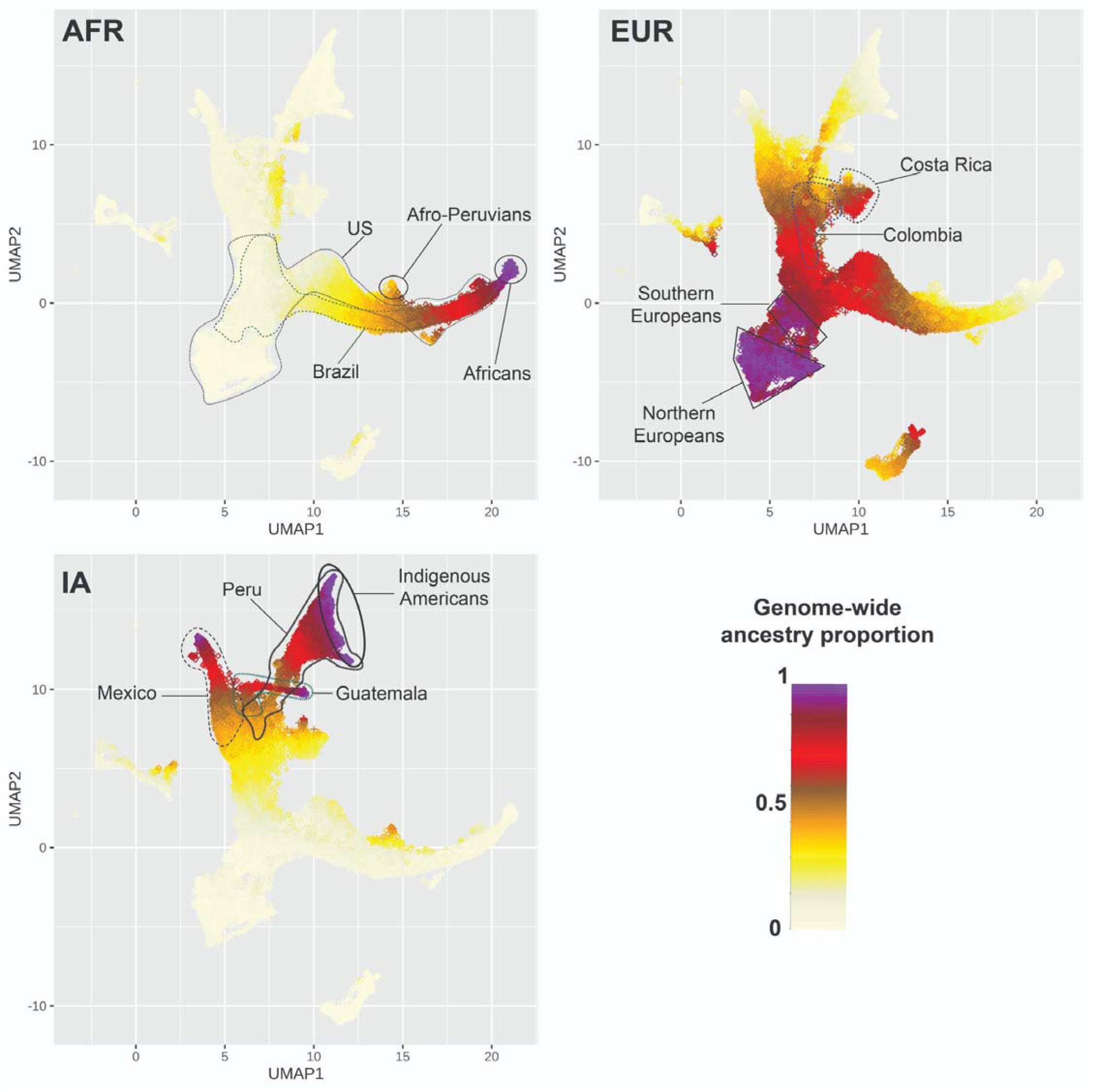
Genome-wide ancestry clines projected on UMAP analysis. Continental ancestry clines based on ancestry proportions inferred by ADMIXTURE for African (AFR), European (EUR) and Indigenous American (IA) ancestries in GLADdb individuals.

**Figure S6A.**
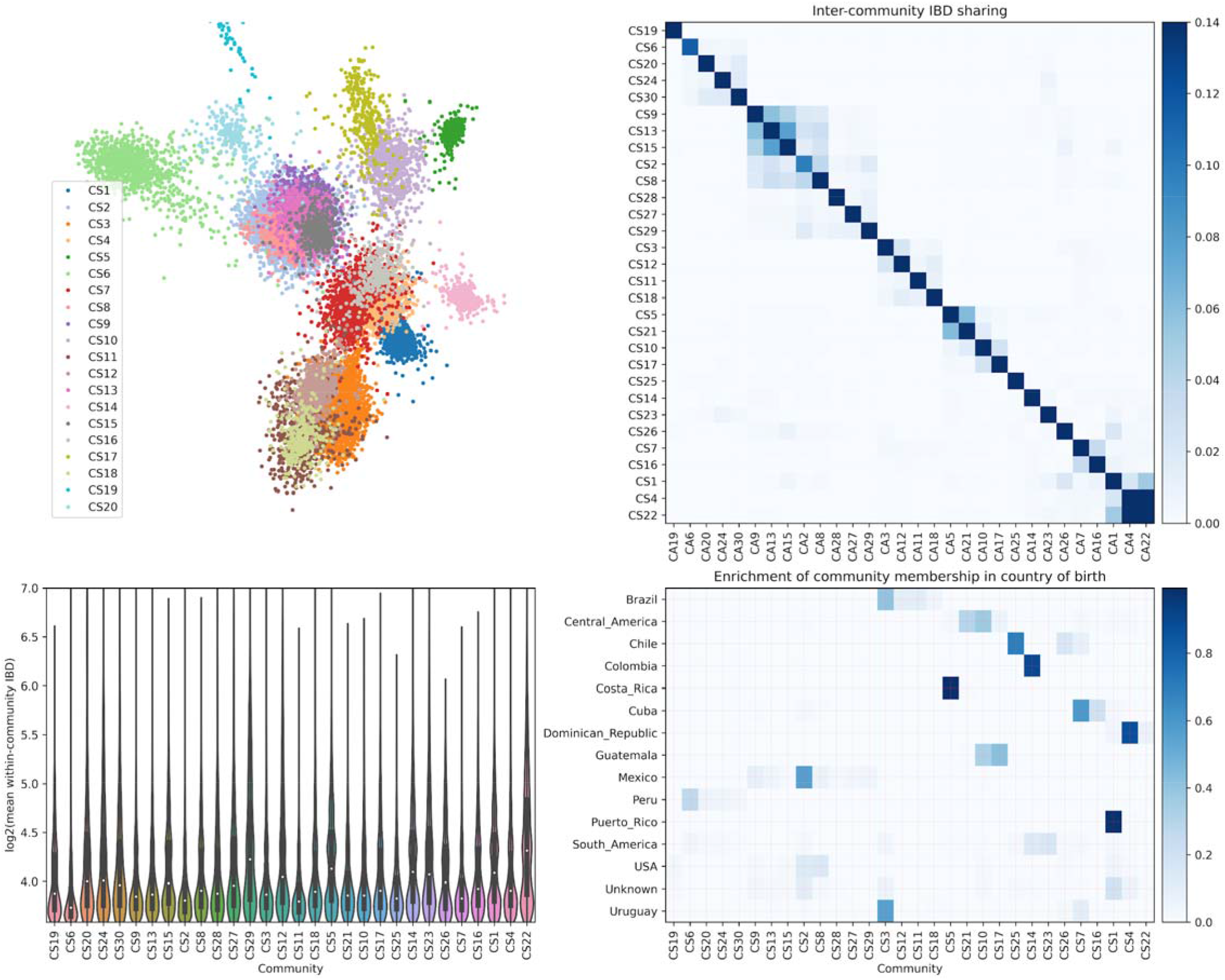
IBD network community detection using IBD segments between 5-9.3cM. This interval was selected to explore the population dynamics before the colonial times. A) Top 20 IBD network communities visualized using Fruchterman-Reingold layout algorithm ^82^. For visualization purposes, only individuals with connections > 30 are included in the layout calculation. The community labels, such as CS1 and CS2, are named according to the IBD version used and the rank of the community sizes, with CS1 representing the largest community when using short IBD segments (5-9.3cM). B) IBD sharing among top 30 inferred communities (ordered by agglomerative clustering; the same order was followed in C and D). C) Distribution of IBD shared among individuals in each community. D) Enrichment of IBD community membership in the country of origin (i.e., proportions of community labels for individuals born in a given country).

**Figure S6B.**
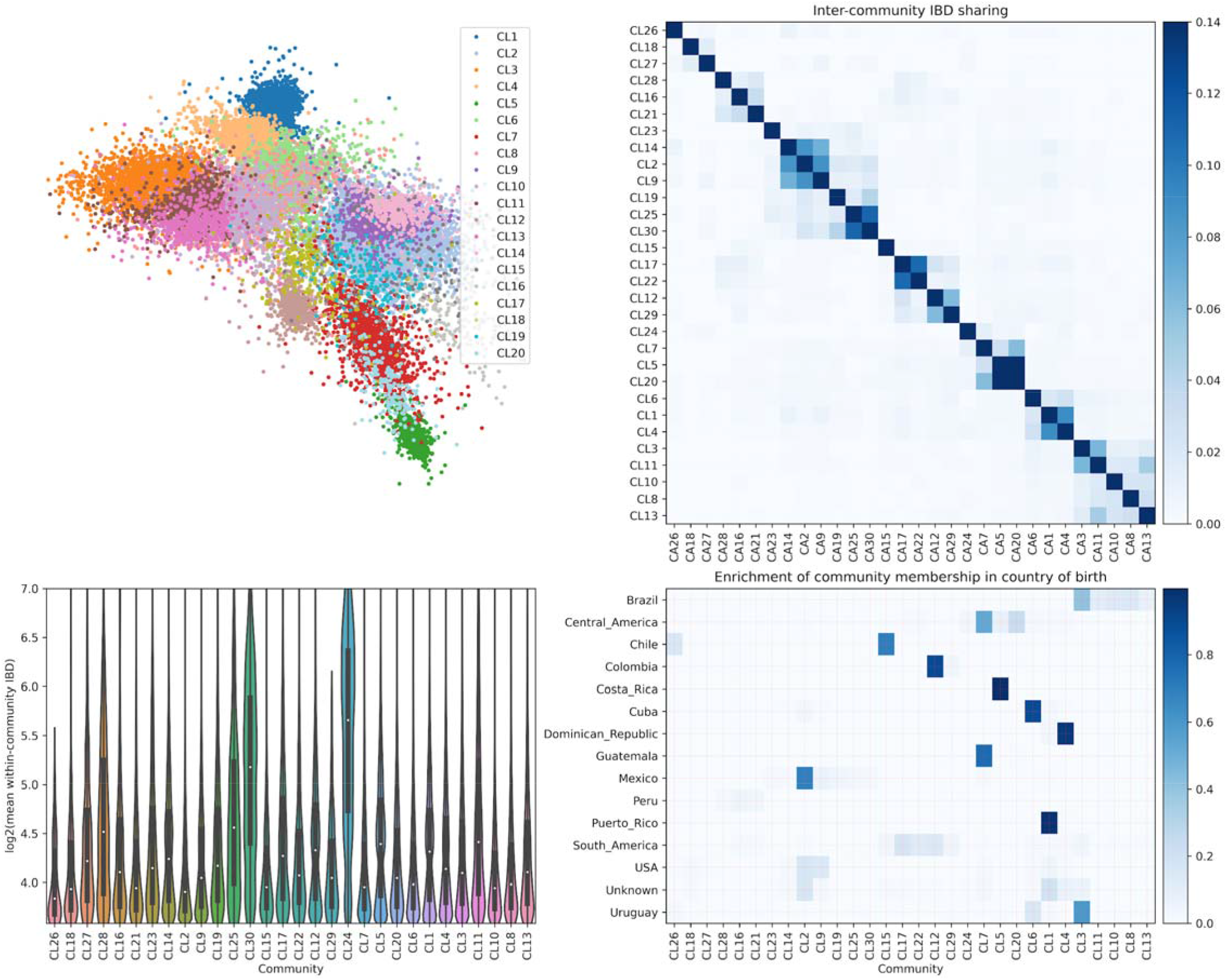
IBD network community detection using IBD segments greater than 9.3cM. This interval was selected to explore the population dynamics after the colonial times. A) Top 20 IBD network communities visualized using Fruchterman-Reingold layout algorithm ^82^. For visualization purposes, only individuals with connections > 30 are included in the layout calculation. The community labels, such as CL1 and CL2, are named according to the IBD version used and the rank of the community sizes, with CL1 representing the largest community when using large IBD segments (> 9.3cM). B) IBD sharing among top 30 inferred communities (ordered by agglomerative clustering; the same order was followed in C and D). C) Distribution of IBD shared among individuals in each community. D) Enrichment of IBD community membership in the country of origin (i.e., proportions of community labels for individuals born in a given country).

**Figure S7A.**
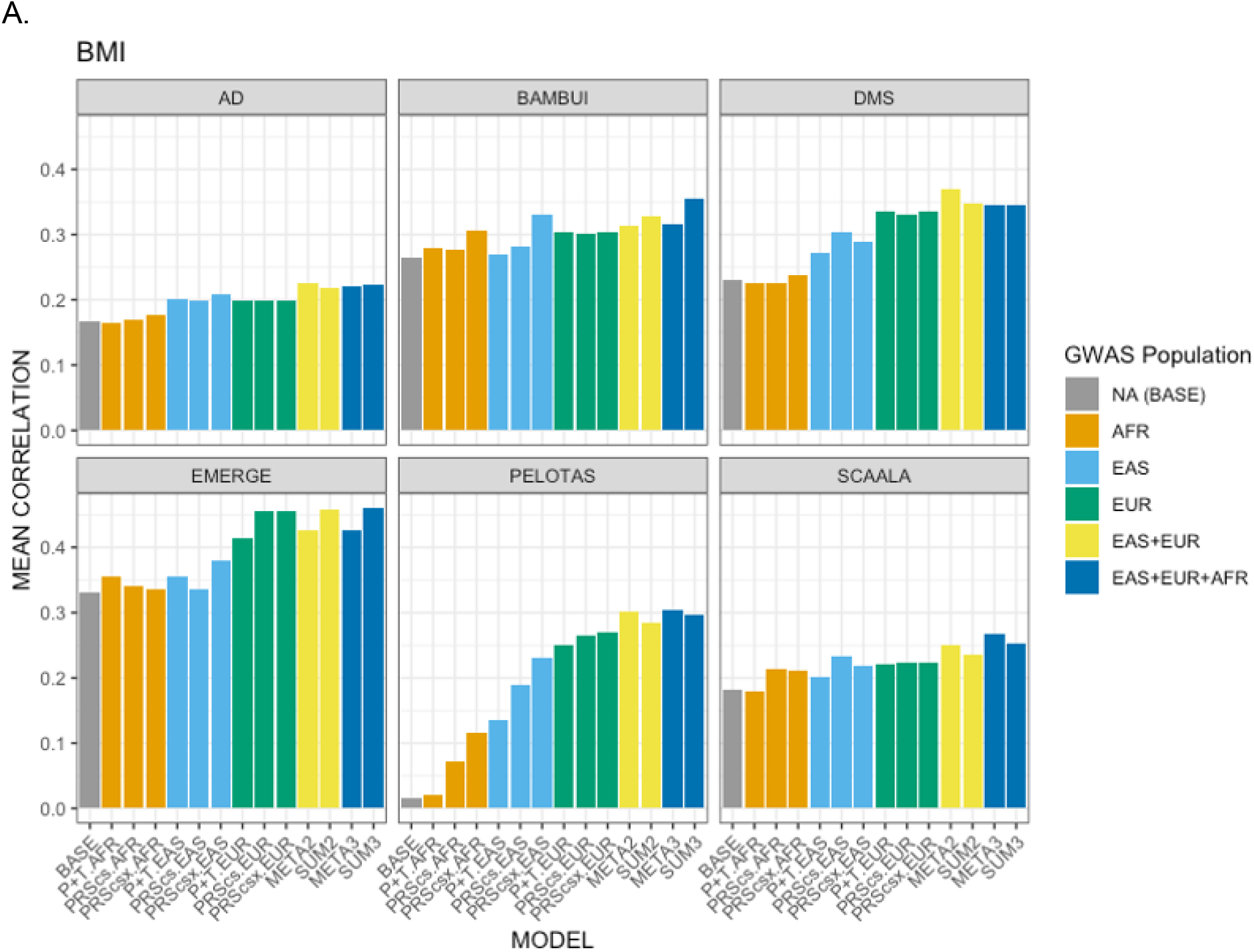
Predictive Performance as measured by the mean correlation of the trait with the prediction. A: Predictive performance for BMI. B: Predictive performance for height. C: Predictive performance for T2D. AD: Columbia University Study of Caribbean Hispanics and Late Onset Alzheimer’s disease (phs000496), DMS: Slim Initiative in Genomic Medicine for the Americas (SIGMA): Diabetes in Mexico Study (phs001388), EMERGE: eMERGE Network Phase III: HRC Imputed Array Data (phs001584), TB: Early Progression to Active Tuberculosis in Peruvians (phs002025), and EPIGEN-Brasil (Bambui, Pelotas, and SCAALA).

**Figure S7B.**
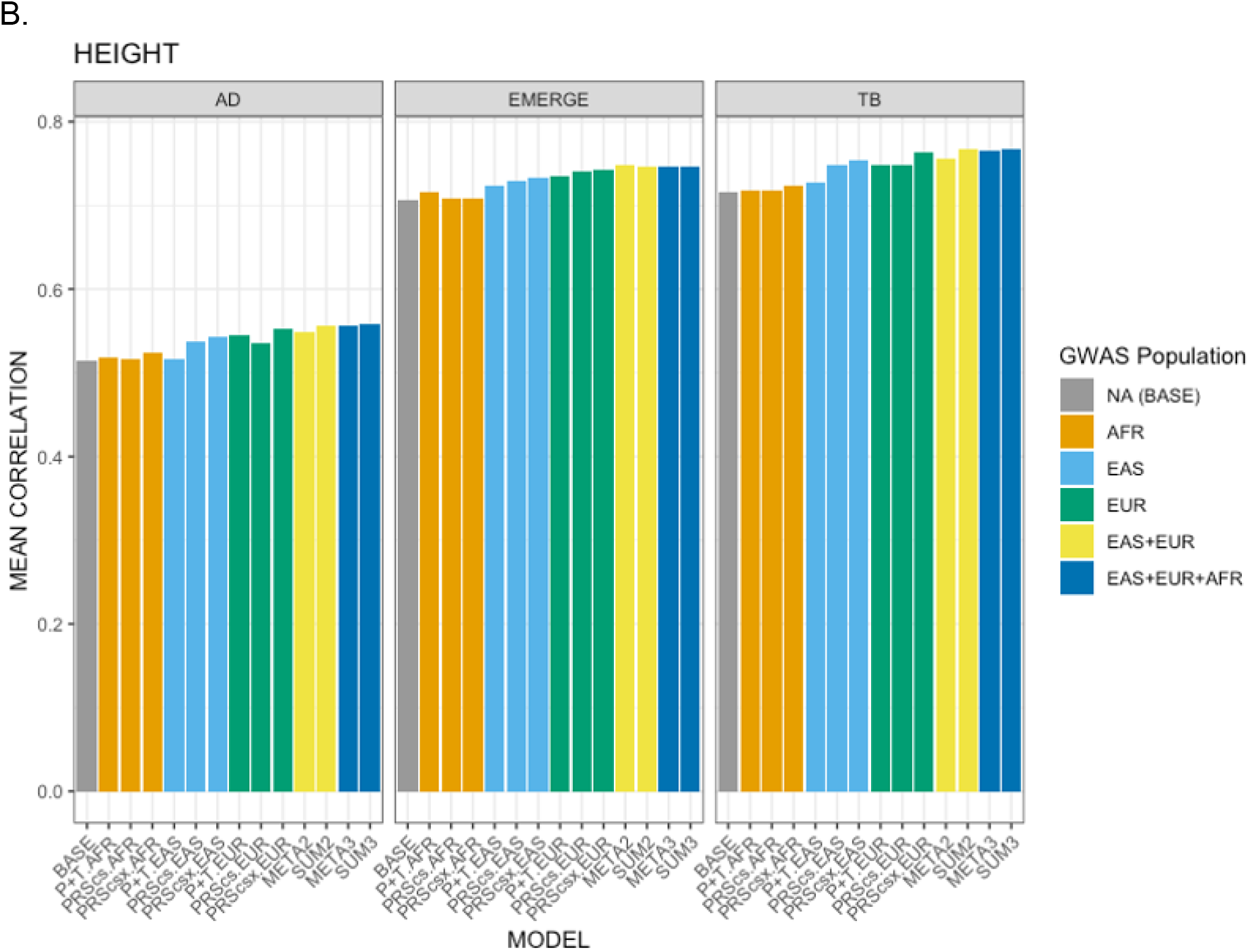
Predictive Performance as measured by the mean correlation of the trait with the prediction. A: Predictive performance for BMI. B: Predictive performance for height. C: Predictive performance for T2D. AD: Columbia University Study of Caribbean Hispanics and Late Onset Alzheimer’s disease (phs000496), DMS: Slim Initiative in Genomic Medicine for the Americas (SIGMA): Diabetes in Mexico Study (phs001388), EMERGE: eMERGE Network Phase III: HRC Imputed Array Data (phs001584), TB: Early Progression to Active Tuberculosis in Peruvians (phs002025), and EPIGEN-Brasil (Bambui, Pelotas, and SCAALA).

**Figure S7C.**
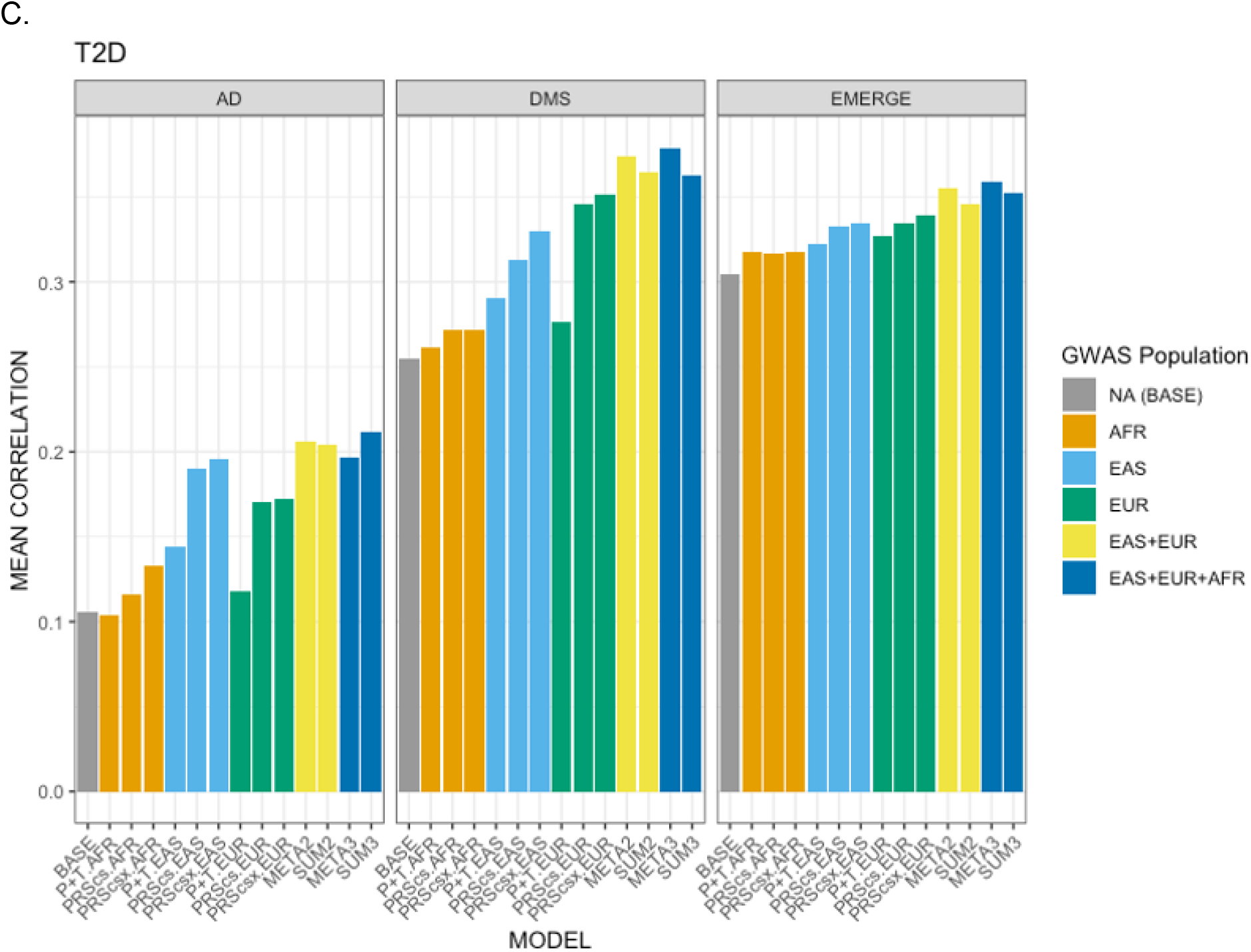
Predictive Performance as measured by the mean correlation of the trait with the prediction. A: Predictive performance for BMI. B: Predictive performance for height. C: Predictive performance for T2D. AD: Columbia University Study of Caribbean Hispanics and Late Onset Alzheimer’s disease (phs000496), DMS: Slim Initiative in Genomic Medicine for the Americas (SIGMA): Diabetes in Mexico Study (phs001388), EMERGE: eMERGE Network Phase III: HRC Imputed Array Data (phs001584), TB: Early Progression to Active Tuberculosis in Peruvians (phs002025), and EPIGEN-Brasil (Bambui, Pelotas, and SCAALA).

**Figure S8.**
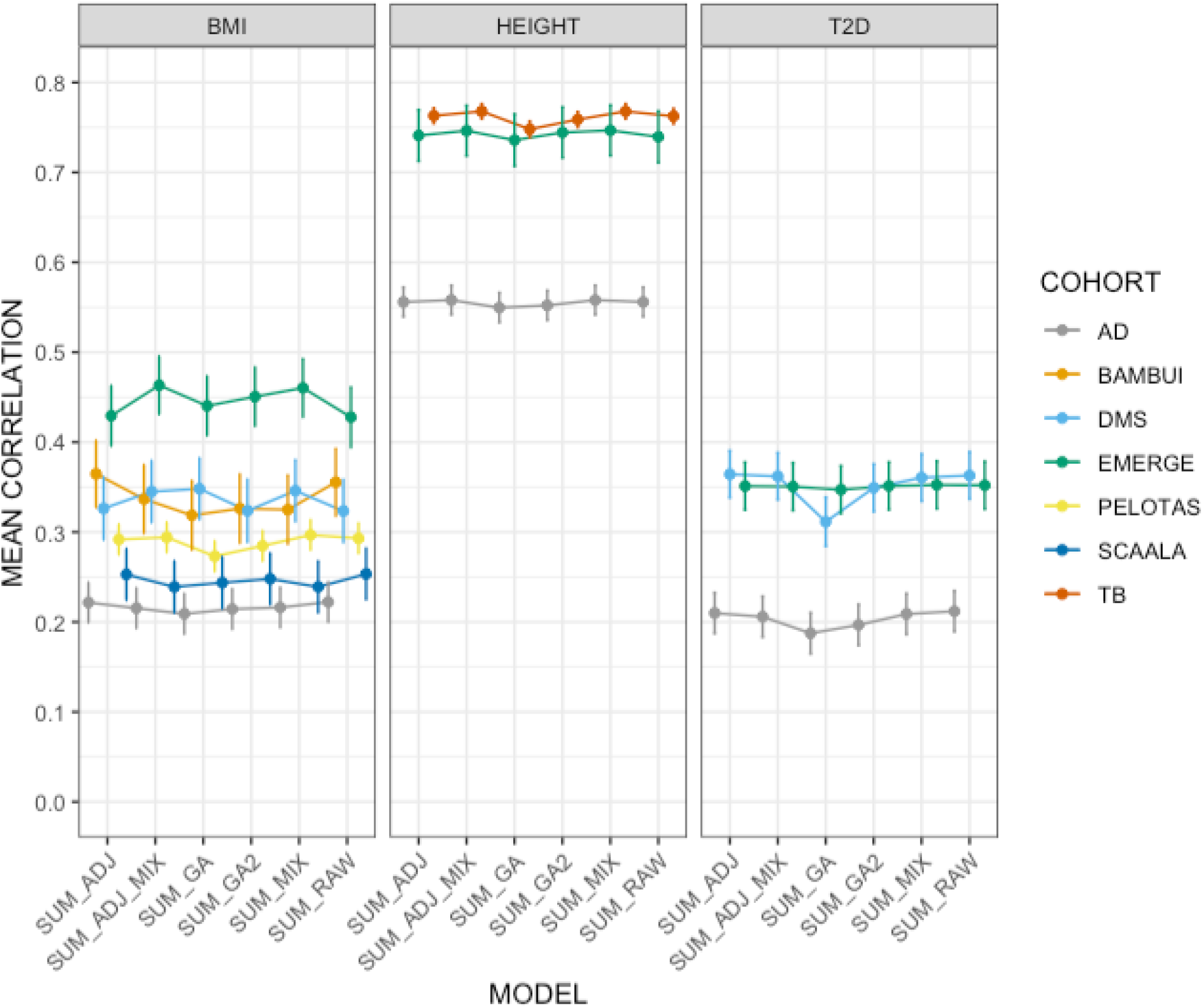
PRS linear combination methods across three traits. Predictive performance of linear combination methods across 3 traits and 7 cohorts. Error bars represent the standard error of the correlation. See methods for model definitions. AD: Columbia University Study of Caribbean Hispanics and Late Onset Alzheimer’s disease (phs000496), DMS: Slim Initiative in Genomic Medicine for the Americas (SIGMA): Diabetes in Mexico Study (phs001388), EMERGE: eMERGE Network Phase III: HRC Imputed Array Data (phs001584), TB: Early Progression to Active Tuberculosis in Peruvians (phs002025), and EPIGEN-Brasil (Bambui, Pelotas, and SCAALA).

**Figure S9.**
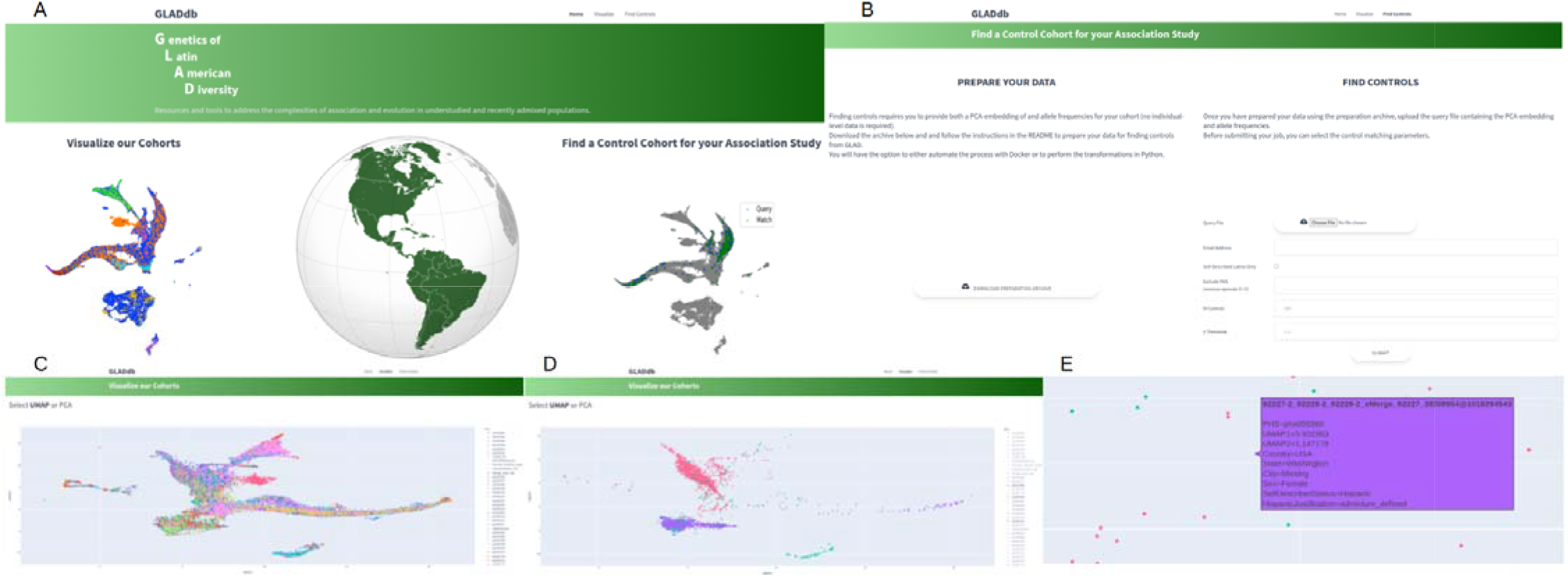
Screenshots from GLADdb website. A) Landing page. B) The control matching page, encompassing data preparation and matching job submission. C-E) Visualization pages, showing respectively all cohorts, selected cohorts, and a zoomed-in view with a highlighted individual.

## Consortia Authorship

### Latin American Research Consortium on the Genetics of Parkinson’s Disease (LARGE-PD)

- Emilia Gatto,
- Grace Letro
- Jorge Luis Orozco
- Carlos Velez-Pardo
- Marlene Jimenez-Del-Rio
- Francisco Lopera
- Patricio Olguin
- Andrew Sobering
- Alex Medina
- Daniel Martinez
- Mayela Rodriguez
- Sarael Alcauter
- Alejandra Medina
- Mario Cornejo-Olivas
- Angel Medina Colque
- Julia Rios Pinto
- Ivan Cornejo Herrera
- Edward Ochoa Valle
- Nicanor Mori Quispe
- Angel Viñuela

### NINDS Stroke Genetics Network (SiGN) Consortium

- Stephen J. Kittner
- Braxton D. Mitchell
- Jordi Jimenez-Conde

### TOPMed Population Genetics Working Group

- Sebastian Zoellner

## Notes

### Competing Interest Statement

The authors have declared no competing interest.

### Summary of Updates

This version now includes abstract versions in Spanish and Brazilian Portuguese in the Main Text. Also, the list of authors has been revised.

