## Supplementary information for "Genetics of Latin American Diversity (GLAD) Project: insights into population genetics and association studies in recently admixed groups in the Americas"

Stephen J. Kittner^a^, Braxton D. Mitchell^b,c^, Jordi Jimenez-Conde^d^.

^a^Department of Neurology, University of Maryland School of Medicine, Baltimore, MD 21201, USA.

^b^Department of Medicine, University of Maryland School of Medicine, Baltimore, MD, 21201,USA.

^c^Geriatrics Research and Education Clinical Center, Baltimore Veterans Administration Medical Center, Baltimore, MD, USA.

^d^Neurovascular Research Group (NEUVAS), Neurology Department, Institut Hospital del Mar d’Investigacio Medica, Universitat Autonoma de Barcelona, Barcelona, Spain

**TOPMed Population Genetics Working Group:**

Sebastian Zoellner^e^

^e^University of Michigan

**Latin American Research Consortium on the Genetics of Parkinson's Disease (LARGE-PD):**

Emilia Gatto^f^, Gracce Letro^g^, Jorge Luis Orozco^h^, Carlos Velez-Pardo^i^, Marlene Jimenez-Del-Rio^i^, Francisco Lopera^i^, Patricio Olguin^j^, Andrew Sobering^k^, Alex Medina^l^, Daniel Martinez^m^, Mayela Rodriguez^n^, Sarael Alcauter^o^, Alejandra Medina^p^, Mario Cornejo-Olivas^q^, Angel Medina Colque^q^, Julia Rios Pinto^q^, Ivan Cornejo Herrera^q^, Edward Ochoa-Valle^q^, Nicanor Mori Quispe^r^, Angel Viñuela^s^.

^f^Department of Neurology, Affiliated University of Buenos Aires, Buenos Aires, Argentina

^g^Pontifícia Universidade Católica de Campinas, Centro de Ciências da Vida, Campinas SP, Brazil

^h^Departamento de Neurología, Fundación Valle del Lili, Cali, Colombia

^i^Neuroscience Research Group, Medical Research Institute, Faculty of Medicine, Universidad de Antioquia (UdeA), Medellín, Colombia

^j^Program in Human Genetics, Institute of Biomedical Sciences, Biomedical Neuroscience Institute, Department of Neuroscience, Facultad de Medicina, Universidad de Chile, Santiago, Chile

^k^Department of Biochemistry, St. George's University School of Medicine, St. George's, Grenada

^l^Hospital de Especialidades San Felipe, Tegucigalpa, Honduras

^m^Tecnologico de Monterrey, Escuela de Medicina y Ciencias de la Salud, Monterrey, MX

^n^Movement Disorders Clinic, National Institute of Neurology and Neurosurgery, Mexico City, MX

^o^Instituto de Neurobiología, UNAM, Campus Juriquilla, Querétaro, México

^p^Laboratorio Internacional de Investigación sobre el Genoma Humano, Universidad Nacional Autónoma de México, Querétaro, México

^q^Neurogenetics Research Center, Instituto Nacional de Ciencias Neurologicas, Lima, Peru

^r^Department of Neurology, Hospital Nacional Daniel A. Carrión, Bellavista, Callao, Peru

^s^Movement Disorders Group, Manatí Medical Center, Neurosciences Institute, Manatí, Puerto Rico.

#### GLADdb

We developed an online portal where investigators both 1) find controls and 2) interact with and visualize GLAD cohorts. In the first use case, investigators can provide summary statistics from their cases and we match and provide summary statistics as controls. As GLADdb samples were ascertained for various phenotypes, options are provided so that samples with known phenotypes are removed from consideration (e.g. in a Parkinson’s Disease (PD) case study any cases with PD are not included as potential controls) as well as the option to remove admixture-defined individuals. No individual-level genotype data is communicated in either direction. For the second use case, we provide a visualization portal wherein users view GLAD samples and cohorts in a variety of embedding spaces (PCA, UMAP). They can visualize samples by cohort, population, gender, etc. to assist them in determining if there is a specific cohort for which they need to apply for individual-level data access. **Figure S9** contains screenshots from the portal. The online portal is hosted on virtual machines, with a separate computer cluster handling the computation required by the matching service.

#### Supplementary information 2 - Study acknowledgments

**NHLBI TOPMed - The Genetic Epidemiology of Asthma in Costa Rica:** This study was supported by NHLBI grants R37 HL066289 and P01 HL132825. We wish to acknowledge the investigators at the Channing Division of Network Medicine at Brigham and Women's Hospital, the investigators at the Hospital Nacional de Niños in San José, Costa Rica 39 and the study subjects and their extended family members who contributed samples and genotypes to the study, and the NIH/NHLBI for its support in making this project possible.

**NHLBI TOPMed - San Antonio Family Heart Study (SAFHS):** Collection of the San Antonio Family Study data was supported in part by National Institutes of Health (NIH) grants R01 HL045522, MH078143, MH078111 and MH083824; and whole genome sequencing of SAFS subjects was supported by U01 DK085524 and R01 HL113323. Sample processing for sequencing was performed in facilities constructed with the support of NIH grant C06 RR020547. ORCID ID: 0000-0001-6250-5723.

**NHLBI TOPMed - Women's Health Initiative (WHI):** The WHI program is funded by the National Heart, Lung, and Blood Institute, National Institutes of Health, U.S. Department of Health and Human Services through contracts HHSN268201600018C, HHSN268201600001C, HHSN268201600002C, HHSN268201600003C, and HHSN268201600004C.

**NHLBI TOPMed - Hispanic Community Health Study - Study of Latinos (HCHS/SOL):** The Hispanic Community Health Study/Study of Latinos is a collaborative study supported by contracts from the National Heart, Lung, and Blood Institute (NHLBI) to the University of North Carolina (HHSN268201300001I / N01-HC-65233), University of Miami (HHSN268201300004I / N01-HC-65234), Albert Einstein College of Medicine (HHSN268201300002I / N01-HC-65235), University of Illinois at Chicago – HHSN268201300003I / N01-HC-65236 Northwestern Univ), and San Diego State University (HHSN268201300005I / N01-HC-65237). The following Institutes/Centers/Offices have contributed to the HCHS/SOL through a transfer of funds to the 47 NHLBI: National Institute on Minority Health and Health Disparities, National Institute on Deafness and Other Communication Disorders, National Institute of Dental and Craniofacial Research, National Institute of Diabetes and Digestive and Kidney Diseases, National Institute of Neurological Disorders and Stroke, NIH Institution-Office of Dietary Supplements.

**NHLBI TOPMed - Multi-Ethnic Study of Atherosclerosis:** MESA and the MESA SHARe projects are conducted and supported by the National Heart, Lung, and Blood Institute (NHLBI) in collaboration with MESA investigators. Support for MESA is provided by contracts 75N92020D00001, HHSN268201500003I, N01-HC-95159, 75N92020D00005, N01-HC-95160, 75N92020D00002, N01-HC-95161, 75N92020D00003, N01-HC-95162, 75N92020D00006, N01-HC-95163, 75N92020D00004, N01-HC-95164, 75N92020D00007, N01-HC-95165, N01-HC-95166, N01-HC-95167, N01-HC-95168, N01-HC-95169, UL1-TR-000040, UL1-TR-001079, UL1-TR-001420, and R01HL105756. Also supported in part by the National Center for Advancing Translational Sciences, CTSI grant UL1TR001881, and the National Institute of Diabetes and Digestive and Kidney Disease Diabetes Research Center (DRC) grant DK063491 to the Southern California Diabetes Endocrinology Research Center. The authors thank the other investigators, the staff, and the participants of the MESA study for their valuable contributions. A full list of participating MESA investigators and institutes can be found at <http://www.mesa-nhlbi.org>.

**NHLBI TOPMed - Severe Asthma Research Program (SARP):** The authors thank the SARP participants, investigators, clinical research staff and data coordinating center. SARP was conducted with the support of the National Institutes of Health (NIH), National Heart, Lung, and Blood Institute (NHLBI) grants R01 HL069116, R01 HL069130, R01 HL069149, R01 HL069155, R01 HL069167, R01 HL069170, R01 HL069174, R01 HL069349, U10 HL109086, U10 HL109146, U10 HL109152, U10 HL109164, U10 HL109168, U10 HL109172, U10 HL109250, and U10 HL109257.

**NHLBI TOPMed - Recipient Epidemiology and Donor Evaluation Study-III Brazil Sickle Cell Disease Cohort (REDS-BSCDC):** The Recipient Epidemiology and Donor Evaluation Study (REDS-III) International Component - Brazil Sickle Cell Disease Cohort Study was funded by National Heart, Lung, and Blood Institute of NIH (Contract No. HHSN268201100007I) and is a collaboration of Blood Systems Research Institute and Research Triangle Institute International in the USA and the University of Sao Paulo, Institute of Tropic Medicine, Fundacao Pro-Sangue, Institute for the Treatment of Childhood Cancer (ITACI), Hemominas, Hemorio, and Hemope in Brazil.

### **NHLBI TOPMed - My Life Our Future (MLOF) Research Repository of Patients with Hemophilia A (Factor VIII Deficiency) or Hemophilia B (Factor IX Deficiency):** My Life Our Future (MLOF) is a project governed by a Steering committee representing Bloodworks Northwest, the American Thrombosis and Hemostasis Network, the National Hemophilia Foundation and Biogen. The project is funded by Biogen. Patients with hemophilia were enrolled at 80 Hemophilia Treatment Centers from across the U.S.

### **NHLBI TOPMed - Boston-Brazil Sickle Cell Disease (SCD) Cohort:** The sickle cell patients included in this cohort were collected by Drs. Igor Domingos, Marcos Andre Bezerra, and Aderson Araujo, and colleagues from their institution, the Hematology and Hemotherapy Foundation of Pernambuco. This study has received partial support from NIH grants R01DK103794 and U01HL117720 (to Vijay G. Sankaran). We are grateful to all study participants for their willingness to be involved in this study.

**NHLBI TOPMed: Children's Health Study (CHS) Integrative Genomics and Environmental Research of Asthma (IGERA):** The Integrative Genomics and Environmental Research of Asthma (IGERA) Study was supported by the National Heart, Lung and Blood Institute (grant # RC2HL101543 -The Asthma BioRepository for Integrative Genomics Research, PI Gilliland/Raby). The Children's Health Study (CHS) was supported by the Southern California Environmental Health Sciences Center (grant P30ES007048); National Institute of Environmental Health Sciences (grants 5P01ES011627, ES021801, ES023262, P01ES009581, P01ES011627, P01ES022845, R01 ES016535, R03ES014046, P50 CA180905, R01HL061768, R01HL076647, R01HL087680 and RC2HL101651), the Environmental Protection Agency (grants RD83544101, R826708, RD831861, and R831845), and the Hastings Foundation.

### **NHLBI TOPMed: Children's Health Study (CHS) Effects of Air Pollution on the Development of Obesity in Children (Meta-AIR):** The Effects of Air Pollution on the Development of Obesity in Children (Meta-AIR) study was supported by the Southern California Children’s Environmental Health Center funded by the National Institute of Environmental Health Sciences (NIEHS) (P01ES022845) and the Environmental Protection Agency (EPA) (RD-83544101–0). The Children's Health Study (CHS) was supported by the Southern California Environmental Health Sciences Center (grant P30ES007048); National Institute of Environmental Health Sciences (grants 5P01ES011627, ES021801, ES023262, P01ES009581, P01ES011627, P01ES022845, R01 ES016535, R03ES014046, P50 CA180905, R01HL061768, R01HL076647, R01HL087680, RC2HL101651 and K99ES027870), the Environmental Protection Agency (grants RD83544101, R826708, RD831861, and R831845), and the Hastings Foundation.

### **NHLBI TOPMed - NHGRI CCDG: The BioMe Biobank at Mount Sinai:** The Mount Sinai BioMe Biobank has been supported by The Andrea and Charles Bronfman Philanthropies and in part by Federal funds from the NHLBI and NHGRI (U01HG00638001; U01HG007417; X01HL134588). We thank all participants in the Mount Sinai Biobank. We also thank all our recruiters who have assisted and continue to assist in data collection and management and are grateful for the computational resources and staff expertise provided by Scientific Computing at the Icahn School of Medicine at Mount Sinai.

### **NHLBI TOPMed - Lung Tissue Research Consortium (LTRC):** This study utilized biological specimens and data provided by the Lung Tissue Research Consortium (LTRC) supported by the National Heart, Lung, and Blood Institute (NHLBI).

### **NHLBI TOPMed - Childhood Asthma Management Program (CAMP):** We thank all subjects for their ongoing participation in this study. We acknowledge the CAMP investigators and research team, supported by NHLBI, for collection of CAMP Genetic Ancillary Study data. All work on data collected from the CAMP Genetic Ancillary Study was conducted at the Channing Laboratory of the Brigham and Women's Hospital under appropriate CAMP policies and human subject's protections. The CAMP Genetics Ancillary Study is supported by U01 HL075419, U01 HL65899, P01 HL083069, R01 HL 086601, R37 HL066289 and T32 HL07427 from the National Heart, Lung and Blood Institute, National Institutes of Health.
